# Dysregulated expression of *Hoxa1* isoforms in hematopoietic stem and progenitor cells causes myelodysplastic syndromes

**DOI:** 10.1101/2023.04.24.538176

**Authors:** ShuhYing Tan, Chacko Joseph, Mohamed Luban Sobah, Alistair M. Chalk, Kelli Schleibs, Samuel C. Lee, Gavin Tjin, Sagrika Chugh, Xiaodong Mo, Clea S. Grace, Jia Q Truong, Monique F. Smeets, Kim L. Rice, Michael W Parker, Ashwin Unnikrishnan, Austin G. Kulasekararaj, Ghulam J. Mufti, Magnus Tobiasson, Eva Hellstrom-Lindberg, John E. Pimanda, Lorraine J. Gudas, Davis J. McCarthy, Jessica K. Holien, Carl R. Walkley, Meaghan Wall, Louise E. Purton

## Abstract

The homeobox gene, *Hoxa1*, has two different isoforms generated by alternative splicing: a full-length homeodomain-containing *Hoxa1* (*Hoxa1-FL*), and a truncated *Hoxa1* (*Hoxa1-T*), that lacks the homeodomain. The effects of the distinct *Hoxa1* isoforms in hematopoiesis have not been investigated. Oncoretroviral studies revealed that Hoxa1-T acts in a dominant negative manner, regulating transcriptionally active Hoxa1. Oncoretroviral overexpression of wildtype *Hoxa1* (*WT-Hoxa1*), which generates both *Hoxa1* isoforms, in murine hematopoietic stem and progenitor cells (HSPCs) perturbed hematopoiesis, resulting in transplantable myelodysplastic syndromes (MDS) in mice. Overexpression of a mutated *Hoxa1* cDNA (*MUT-Hoxa1*) that generates *Hoxa1-FL*, but not *Hoxa1-T*, led to a more severe MDS that transformed to secondary acute myeloid leukemia (sAML). DNA damage repair pathways were downregulated in *Hoxa1*-overexpressing hematopoietic progenitor cells, accompanied by increased γH2AX foci. *In silico* analyses revealed that CD34+ cells from approximately 50% of patients with MDS had elevated *HOXA1-FL* expression. Conditional knock-in *WT-Hoxa1* and *MUT-Hoxa1* mice were generated and had features of pre-MDS, developing altered hematopoiesis within 4 months of *Hoxa1* isoform overexpression in HSPCs. HSPCs were significantly reduced in all knock-in mice, accompanied by significantly increased apoptosis in *WT-Hoxa1* HSPCs. Healthy wildtype recipients transplanted with bone marrow cells from *Hoxa1* knock-in mice developed trilineage MDS, with *Hoxa1* isoform and gene dosage dependent phenotypes. Collectively our data identify a role for HOXA1 in the pathogenesis of MDS. Our *Hoxa1* mouse models capture different stages of progression of disease from pre-MDS to MDS to sAML and provide novel, clinically relevant tools to study MDS.

**Key points:** *HOXA1* is upregulated in approximately 50% of MDS patient CD34+ BM cells, highlighting a potential role for *HOXA1* in the pathogenesis of MDS.

Dysregulated expression of *Hoxa1* isoforms in murine hematopoietic stem and progenitor cells predisposes mice to pre-MDS and MDS.

## Introduction

Homeobox (HOX) proteins are a group of highly conserved transcription factors, defined by the presence of a 60 amino acid DNA-binding domain termed the homeodomain. The target selectivity of HOX proteins is enhanced by interactions with their transcriptional cofactors; Pre-B cell leukemia transcription factor (PBX) proteins and myeloid ecotropic viral integration site (MEIS) proteins. *HOX* genes are arranged in clusters and the HOX proteins from the 3’ end of the cluster form heterodimers with PBX proteins whereas HOX proteins from the 5’ end interact with MEIS proteins ^1–3^. The existence of HOX spliced variants that lack homeodomains have been described for Hoxb6, Hoxa9 and Hoxa1 in mice and their human counterparts HOXB6, HOXA9 and HOXA1 ^4–8^. The functions of these truncated HOX proteins remain largely unknown ^6,9^.

*Hoxa1* is one of the earliest *Hox* genes to be expressed and is essential for normal embryogenesis ^10^. *Hoxa1* null mice display hindbrain segmentation and patterning defects resulting in embryonic lethality ^11,12^. *Hoxa* and *Hoxb* cluster genes are expressed in normal murine adult hematopoietic cells, with the highest level of expression restricted to hematopoietic stem cells (HSCs) ^13,14^. Increased levels of *HOX* mRNA or translocations involving *HOX* genes have been reported in human hematological malignancies ^15–18^. The *Hoxa* gene cluster has also been shown to be essential for murine HSPCs ^19^.

The wild type *Hoxa1* (*WT-Hoxa1*) gene generates two distinct transcripts by alternative splicing: full length *Hoxa1* (*Hoxa1-993*, hereafter identified as *Hoxa1-FL*) and truncated *Hoxa1* (*Hoxa1-399*, referred to here as *Hoxa1-T*) ^5^. *Hoxa1-FL* encodes a protein of 331aa containing a DNA homeobox-binding domain and regulates gene transcription by binding to DNA and its cofactor, PBX1. In contrast, *Hoxa1-T* encodes a 133aa protein due to the early appearance of a stop codon caused by a reading frame shift from alternative splicing within exon 1 of *Hoxa1-FL*. HOXA1-T and HOXA1-FL are identical in their first 114aa, however, HOXA1-T does not contain the homeobox domain, therefore lacks the ability to bind to DNA. The human *HOXA1* gene shares 90% sequence homology with its murine counterpart and likewise generates different isoforms through alternative splicing ^8^.

Myelodysplastic syndromes (MDS) are clonal myeloid malignancies characterized by ineffective hematopoiesis, resulting in peripheral blood cytopenia/s accompanied by morphological evidence of dysplasia in one or more lineages, and carry a heightened risk of transformation to secondary AML (sAML) ^20^. The MDS-initiating cell has been postulated to arise from CD34+ HSPCs, which accumulate co-operating genetic and epigenetic mutations to drive disease progression ^21^.

MDS is a very heterogeneous disease. There are over 50 recurrently mutated genes in MDS, and these somatic mutations are apparent in 80-90% of patients, including those previously found to have a normal karyotype by conventional cytogenetic analysis ^22,23^. Despite improved knowledge of the disease there are limited therapeutic options for MDS patients ^24^. Furthermore, many of the models available for the study of MDS either do not completely mimic the human disease or represent less than 10% of MDS patients ^21,25^.

It has been recognized that MDS can be preceded by clinical conditions referred to as “pre-MDS”^26^. Two of these conditions: clonal hematopoiesis of indeterminate potential (CHIP) and idiopathic dysplasia of unknown significance (IDUS) are associated with normal peripheral blood counts. Another two have persistent cytopenias, being clonal cytopenia of undetermined significance (CCUS) and idiopathic cytopenia of unknown significance (ICUS)^26^. Known somatic mutations are detected in CHIP and CCUS patient cells but not in those from IDUS and ICUS^26^. Not all pre-MDS patients develop MDS and it is currently not clear how pre-MDS evolves into MDS ^26,27^.

Here we show that altered expression of the *Hoxa1* isoforms in murine HSPCs resulted in distinct, novel mouse models of pre-MDS and MDS. We confirmed that *HOXA1* was upregulated in CD34^+^ BM cells from approximately 50% of patients with MDS. Collectively our data suggest a role for HOXA1 in the pathogenesis of MDS.

## Methods

### Animal models

All animal experiments were approved by the St. Vincent’s Hospital Animal Ethics Committee, Melbourne and performed in compliance with the Australian Code of Practice for the Care and Use of Animals for Scientific Purposes (2013).

### Retroviral overexpression models

cDNAs of *WT-Hoxa1*, *MUT-Hoxa1* and *Hoxa1-T* were cloned into the MND-X-IRES-*eGFP* (MXIE) vector ^28^. *MUT-Hoxa1* was generated by site-directed mutagenesis using standard conditions. Retroviral vectors, transductions and cell cultures were generated and performed using methods previously described ^29^.

Transplantation studies were performed using 8-week-old mice purchased from the Animal Resource Centre (WA, Australia). Recipients were lethally irradiated (10Gy total dose, two split doses given 3 hours apart) using a Gammacell 40 Exactor (Best Theratronics, ON, Canada) prior to transplantation. 1 to 5 x 10^6^ unsorted BM cells from B6.SJL-Ptprca^Pep3b/BoyJArc^ (B6.SJL Ptprca mice) were transplanted into primary recipient C57BL/6 mice immediately after transduction, without selection for GFP+ cells. Each primary recipient within each group was transplanted with an aliquot of separately-transduced cells. Four months after the primary transplants, 2 x 10^6^ BM cells from primary recipients of *WT-Hoxa1*, *MUT-Hoxa1* or MXIE transduced cells were transplanted into irradiated secondary recipients. All studies were repeated, and the results were pooled within vector groups.

### Conditional knock-in models

Conditional knock-in mice were generated by Taconic Biosciences (Köln, Germany). *WT- Hoxa1* or *MUT-Hoxa1* was inserted into exon 1 of the *Rosa26* (*R26*) gene, with a stop codon flanked by loxP sites placed immediately upstream. The mice were bred to *hScl*-Cre^ERT^ C57BL/6 mice ^30^, and the offspring were then backcrossed to produce congenic strains expressing the knock-in in a heterozygous (*WT-Hoxa1^ki/+^* or *MUT-Hoxa1^ki/+^*) or homozygous manner (*WT-Hoxa1^ki/ki^* or *MUT-Hoxa1^ki/ki^*). Tamoxifen-treated Cre^+^ littermate mice that lacked the knock-in were used as controls for all experiments.

Mice were placed on tamoxifen-containing chow (400mg/kg tamoxifen citrate; Specialty Feeds, Perth WA) for 4 weeks commencing at 4 weeks of age. At 4 months after induction of the knock-ins, mice were euthanized for complete hematological analyses, and BM cells were used for transplantation studies.

Prior to transplantation, 8-week-old B6.SJL-Ptprca^Pep3b/BoyJArc^ male recipients were lethally irradiated. Whole BM (2 x 10^6^ cells) was transplanted into irradiated recipients, and mice were monitored long-term for the development of MDS. All studies were repeated, and the results were pooled within knock-in groups.

### Flow cytometry analysis of hematopoietic cells

Cells were counted, stained and analyzed by flow cytometry as previously described ^31^.

### Gene Expression Analysis

Gene expression data for 183 MDS CD34^+^ cells and 17 controls ^32^ were obtained from Gene Expression Omnibus microarray database (GEO accession number GSE19429). RNA-seq data for 44 MDS and 23 controls ^33^ were likewise obtained (GEO accession number GSE111085).

For murine gene profiling studies, BM GFP+ CMPs and MEPs were sorted from primary recipient mice at 16 weeks post-transplant (n= 4 independent transduction groups/vector, with 4 mice per group). Total RNA was extracted using the RNeasy Micro kit (Qiagen). RNA was hybridized to the Affymetrix Mouse 1.1 ST genechip at the Ramaciotti Centre for Gene Function Analysis at the University of New South Wales, Australia.

Raw data files are available at GEO using accession number GSE62853 for MEPs and GSE227775 for CMPs. Details of microarray analyses and other relevant methods are provided in Supplemental Information.

### Statistical analyses

One-way analysis of variance (ANOVA) or two-way ANOVA followed by post-hoc testing were used for multiple comparisons. The paired or unpaired two-tailed Student’s t-tests were used for other statistical comparisons where indicated. Statistical analysis was performed using Prism 10 software (GraphPad), including assessment of normality, with p<0.05 considered significant.

## Results

### *Hoxa1* isoforms have distinct effects on immature murine hematopoietic cells

We generated vectors containing cDNA encoding *WT-Hoxa1* (which produces both *Hoxa1-FL* and *Hoxa1-T*, Figure 1A) or only *Hoxa1-T* (Figure 1B). A mutated *Hoxa1* (*MUT-Hoxa1*) cDNA was generated by altering the acceptor splice site from AGC (serine) to TCT (serine), conserving the protein sequence of HOXA1-FL but preventing the *Hoxa1-FL* mRNA from undergoing splicing to generate *Hoxa1-T* (Figure 1C). *Hoxa1-T* mRNA was not detected in stable cell lines overexpressing *MUT-Hoxa1* (Figure 1D).

**Figure 1.**
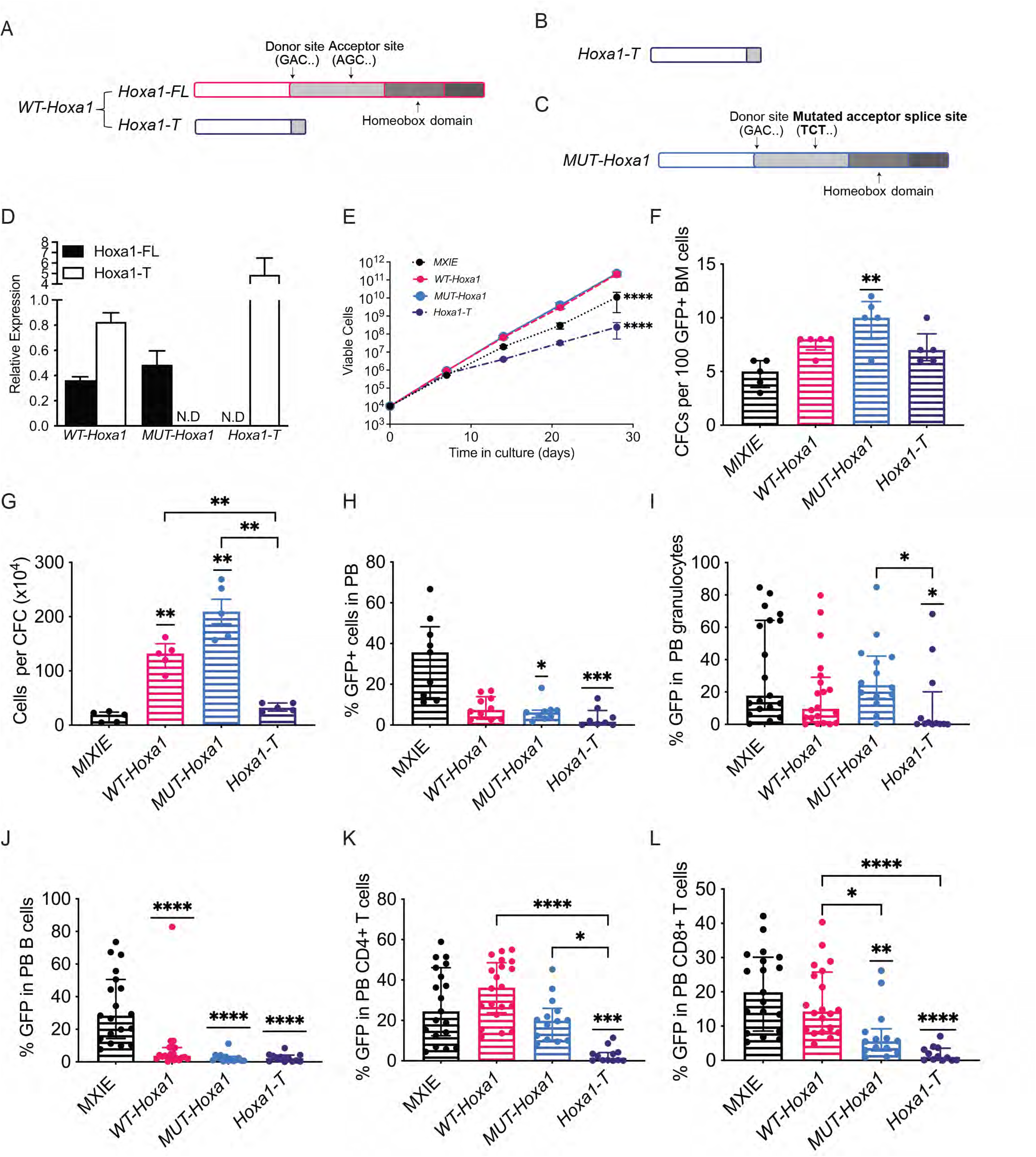
*Hoxa1* isoforms have distinct roles in hematopoiesis. **(A)** Schematic representation of *WT-Hoxa1* and (**B)**, *Hoxa1-T. WT-Hoxa1* generates two isoforms: *Hoxa1-FL*, which contains the DNA-binding homeobox domain and *Hoxa1-T*, which lacks the homeobox domain. Donor and acceptor splice sites are marked on the *Hoxa1-FL* isoform. Regions complementary between *Hoxa1-FL* and *Hoxa1-T* are shown in light grey. (**C**) The mutated form of *Hoxa1* (*MUT-Hoxa1*) generates normal Hoxa1-FL protein but cannot undergo alternative splicing to generate Hoxa1-T. (**D)** Relative expression levels of *Hoxa1-FL* and *Hoxa1-T* in *WT-Hoxa1, MUT-Hoxa1 and Hoxa1-T* overexpressing GP+E-86 packaging cells analyzed by QPCR. (ND: not detected). (**E**) Cumulative cell counts of immature BM cells transduced with MXIE (empty vector) control, *WT-Hoxa1, MUT-Hoxa1* or *Hoxa1*-T cultured *in vitro* (shown is one experiment, representative of four separate experiments). **(F)** Numbers of CFCs after 12 days and (**G)**, Number of cells per CFC produced from 100 LKS^+^ cells transduced with *WT-Hoxa1*, *MUT-Hoxa1*, *Hoxa1-T* or MXIE (n=5). PB parameters of donor cells of *WT-Hoxa1*, *MUT-Hoxa1, Hoxa1-T* and MXIE primary transplant recipients at 16 weeks: **(H)**, Percentage of GFP+ cells in: donor cells; (**I)**, Granulocytes; **(J)**, B lymphocytes; **(K)**, CD4^+^ T lymphocytes; **(L)**, CD8^+^ T lymphocytes (n=12-20). Data in F, H-L are shown as median ± IQR, data shown in G are mean ± SEM; *p<0.05; ** p<0.01; *** p<0.001, ****p<0.0001 vs MXIE control (underlined asterisks above the *Hoxa1* genotype) or between groups designated by the bars. E was analyzed using two-way ANOVA with multiple comparisons, the asterisks indicate significant differences compared to both *WT-Hoxa1* and *MUT-Hoxa1*-overexpressing cells. F-L were analyzed using one-way ANOVA with multiple comparisons.

Overexpression of *WT-Hoxa1* or *MUT-Hoxa1* in immature murine BM cells resulted in extensive *in vitro* proliferation, with >50-fold increases in cell numbers weekly, producing significantly more cells than MXIE (empty vector control) and *Hoxa1-T*-overexpressing cells (Figure 1E). There were significantly increased numbers of colony-forming cells (CFCs) from *MUT-Hoxa1*-overexpressing GFP+ sorted lineage negative, c-kit+ Sca-1+ (LKS^+^) cells (Figure 1F). Significantly increased numbers of cells per colony were observed in both *WT-Hoxa1-* and *MUT-Hoxa1*-overexpressing CFCs, with *MUT-Hoxa1*-overexpressing cells having a 1.6-fold increase in cells per colony compared to *WT-Hoxa1*-overexpressing cells (Figure 1G). *Hoxa1-T* overexpression did not alter the CFC potential of the cells (Figure 1F-G).

### *WT-Hoxa1-* and *MUT-Hoxa1*-overexpression perturbs hematopoiesis with apparent cytopenias demonstrated *in vivo*

Cohorts of lethally irradiated mice were transplanted with BM cells overexpressing either MXIE control, *WT-Hoxa1*, *MUT-Hoxa1* or *Hoxa1-T*. Hereafter, we refer to mice as MXIE (controls), *WT-Hoxa1*, *MUT-Hoxa1 and Hoxa1-T* respectively.

At 16 weeks post-transplant (BMT), *WT-Hoxa1*, *MUT-Hoxa1* and *Hoxa1-T* demonstrated reduced proportions of GFP+ cells in the peripheral blood (PB) compared to MXIE (Figure 1H). *MUT-Hoxa1* had significantly lower numbers of leukocytes and platelets and *Hoxa1-T* had significantly reduced platelets compared to MXIE (Table S1).

*WT-Hoxa1*, *MUT-Hoxa1* and *Hoxa1-T* showed altered repopulation of PB leukocyte subsets (measured as % GFP+ cells detected in each cell type). There was significantly reduced repopulation of granulocytes in *Hoxa1-T* versus *MUT-Hoxa1* (Figure 1I). All three groups displayed significantly reduced repopulation of B lymphocytes when compared to MXIE (Figure 1J) and *Hoxa1-T* had significantly reduced repopulation of CD4^+^ T lymphocytes compared to all other groups (Figure 1K). *Hoxa1-T* and *MUT-Hoxa1* had poor repopulation of CD8^+^ T lymphocytes compared to both MXIE and *WT-Hoxa1* (Figure 1L).

Bone marrow cellularities and the GFP+ BM cells in the MXIE and *Hoxa1* recipient mice were not significantly different (Figure 2A-B). The repopulation of immature and mature BM B lymphocytes were dramatically reduced in both *WT-Hoxa1* and *MUT-Hoxa1* (Figure 2C-D). There were no differences in the repopulation of immature or mature granulocytes (Figure 2E-F).

**Figure 2.**
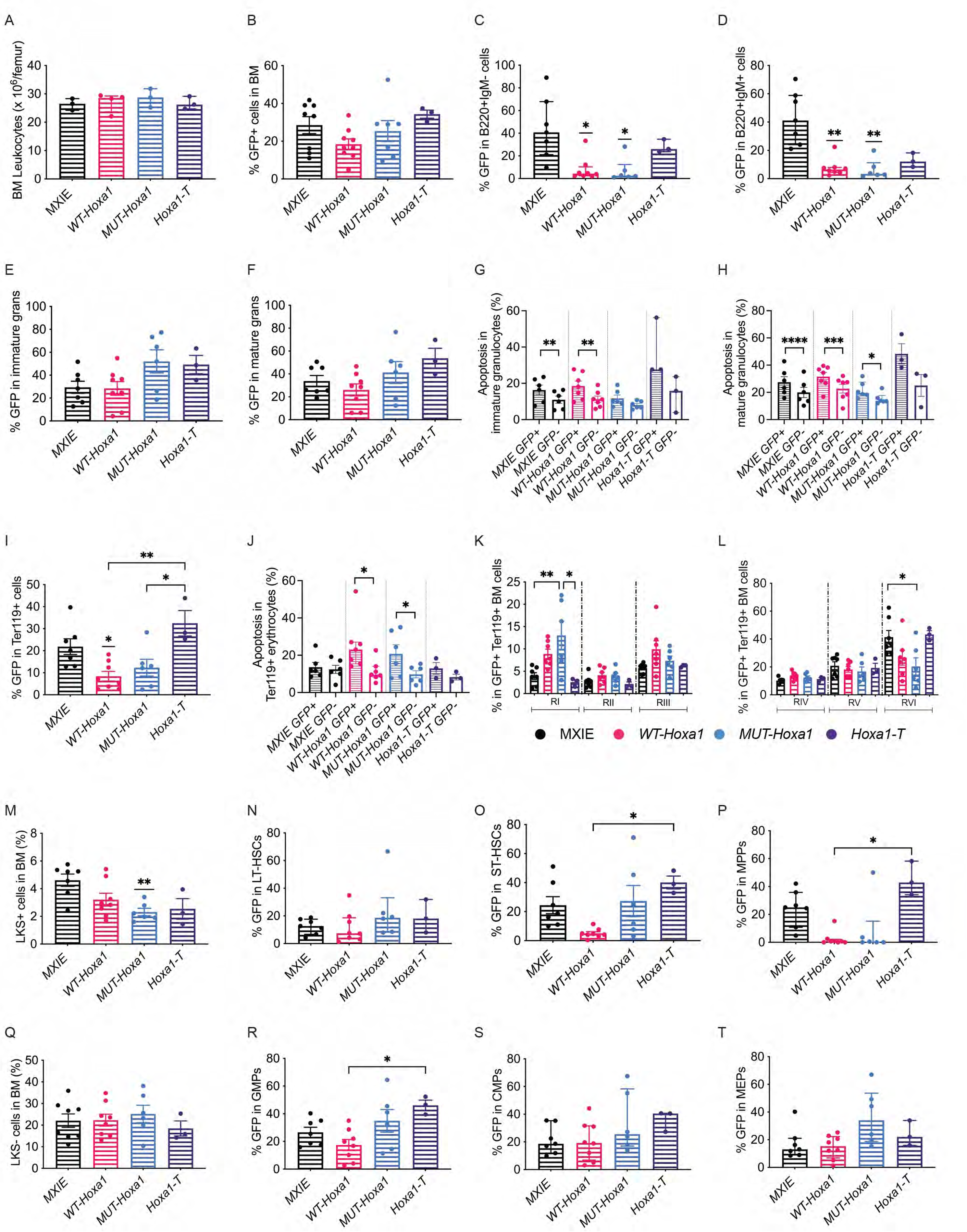
*WT-Hoxa1* or *MUT-Hoxa1* overexpression results in reduced HSPCs and anemia with associated ineffective erythropoiesis. Shown are data from BM harvested from *WT-Hoxa1*, *MUT-Hoxa1, Hoxa1-T* and MXIE recipients at 16 weeks post-transplant: (**A)**, Leukocyte cellularity; (**B)**, Percentage of GFP+ cells in BM leukocytes; (**C)**, Percentage of GFP+ cells in immature (B220^+^IgM^-^) B lymphocytes, (**D)**, Percentage of GFP+ cells in maturing (B220^+^IgM^+^) B lymphocytes. (**E)**, Percentage of GFP^+^ cells in immature (Gr1^lo^CD11b^+^) granulocytes and (**F)**, mature (Gr1^hi^CD11b^+^) granulocytes (n=7-8 recipients). Percentage of apoptotic cells (Annexin V^+^ 7-AAD^-^) in (**G)**, immature and (**H)**, mature BM granulocytes (n=3-7 recipients). Percentage of (**I)**, GFP+ cells in Ter119^+^ erythroid cells; (**J)**, proportions of apoptotic cells (Annexin V^+^ 7-AAD^-^) in erythroid cells (n=3-7 recipients). Percentage of erythrocyte subsets in GFP+ Ter119^+^ BM cells: **(K)** RI proerythroblasts, RII basophilic erythroblasts, RIII polychromatic erythroblasts; (**L**) RIV orthochromatic erythroblasts, RV reticulocytes, RVI mature erythrocytes. **(M)** percentages of LKS^+^ cells in lineage negative BM and the percentages of GFP+ cells in: **(N)**, LT-HSCs; (**O)**, ST-HSCs; (**P)**, MPPs. Percentage of (**Q)**, LKS^-^ cells in lineage negative BM and the percentages of GFP+ cells in: (**R)**, GMP; (**S)**, CMP and (**T)**, MEP (n=3-8). All data are shown as mean ± SEM except C-D, N, P, S and T, which show median ± IQR; *p<0.05; ** p<0.01; *** p<0.001; ****p<0.0001 vs. MXIE control (underlined asterisks above the *Hoxa1* genotype) or between groups designated by the bars. The paired student’s T-test was used for analyses of GFP+ vs GFP- cells in G, H and J; all other data were analyzed by one-way ANOVA with multiple comparisons. Data are pooled from separate experiments.

The MXIE and *WT-Hoxa1* GFP+ immature and mature granulocytes had significantly increased apoptosis compared to GFP- granulocytes within the same mice (Figure 2G-H). There were no differences in apoptotic cells in the immature granulocytes in *MUT-Hoxa1* BM (Figure 2G), however, apoptosis in the GFP+ *MUT-Hoxa1* mature granulocytes was significantly increased (Figure 2H).

*WT-Hoxa1* and *MUT-Hoxa1* mice had significantly reduced proportions of BM erythroid cells compared to both MXIE and *Hoxa1-T*, accompanied by increased apoptosis in GFP+ erythrocytes (Figure 2I-J). Analysis of erythroid subsets^34^ within the GFP+ Ter119+ erythroid cells revealed that *MUT-Hoxa1* BM had increased proportions of RI proerythroblasts and significantly reduced proportions of RVI mature erythrocytes compared to MXIE BM (Figure 2K-L).

### *WT-Hoxa1* and *MUT-Hoxa1* mice have altered numbers of HSPCs

There were significant reductions in HSPC-containing LKS^+^ cells in *MUT-Hoxa1* mice (Figure 2M). The proportions of GFP+ cells in long-term repopulating HSCs (LT-HSCs) were similar across the groups (Figure 2N). There were significant reductions in GFP+ cells in short-term repopulating HSCs (ST-HSCs, Figure 2O) and multipotent progenitors (MPPs, Figure 2P) in *WT-Hoxa1* compared to *Hoxa1-T*.

There were no changes in the proportions of the myeloid progenitor cells (LKS^-^) (Figure 2Q). Aside from significantly increased proportions of GFP+ cells in granulocyte macrophage progenitors (GMPs) of *Hoxa1-T* compared to *WT-Hoxa1* recipients (Figure 2R), no changes were observed in the repopulation of common myeloid progenitors (CMPs), GMPs or megakaryocyte erythroid progenitors (MEPs) across the recipient groups (Figure 2S-T).

### *MUT-Hoxa1* recipients develop sAML

Secondary transplants of BM obtained from MXIE, *WT-Hoxa1* and *MUT-Hoxa1* recipients were performed at 16 weeks post-BMT. With the exception of two cohorts of mice that developed sAML due to a co-operating *Meis1* integration site (Table S2), none of the recipients of *WT-Hoxa1* BM developed sAML (Figure 3A), consistent with a previous report ^35^. However, *WT-Hoxa1* mice developed mortalities associated with low blood counts. Analysis of surviving mice at 6 months post-BMT revealed that there were no significant differences in the proportions of GFP+ donor cells in the PB of *WT-Hoxa1* compared to MXIE controls (Figure 3B). In contrast, they had significantly reduced PB leukocyte counts (Figure 3C) and anemia (Figure 3D-E).

**Figure 3.**
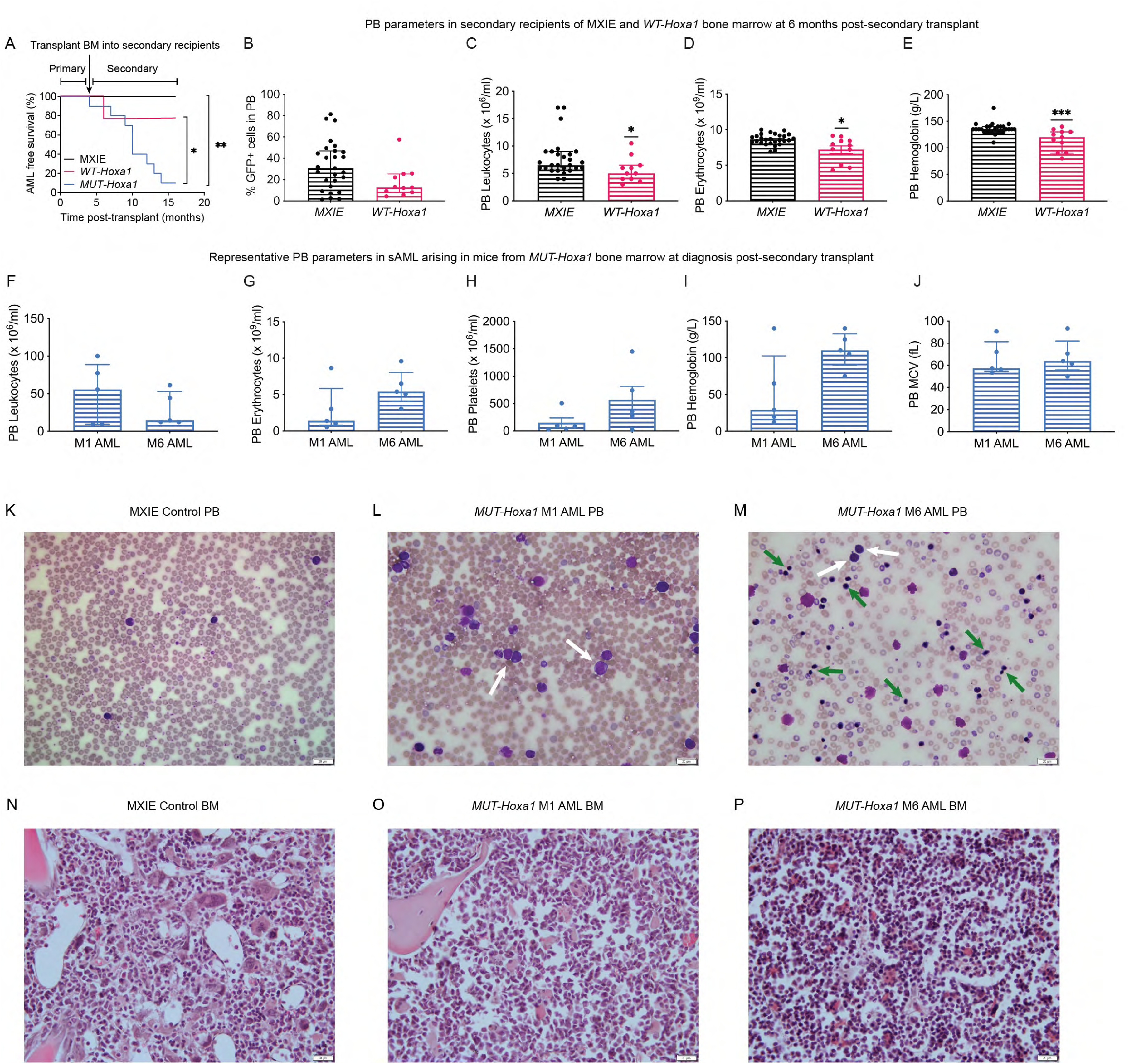
*MUT-Hoxa1* spontaneously progresses to sAML. Shown are: (**A)** Kaplan-Meier survival curve for recipients of *MUT-Hoxa1*, *WT-Hoxa1* and *MXIE* overexpressing BM (n=10 independent transductions, each with 4 recipients per group, only the first mouse to develop sAML is shown per cohort, however, all mice in each cohort developed sAML within 2 weeks of the first mouse). Results were analyzed with a log-rank (Mantel–Cox) test. *p<0.05, **p<0.01 for the groups designated by the bars. PB parameters of secondary recipients of MXIE and *WT-Hoxa1* BM at 6 months post-secondary transplant: (**B)** Percentage of GFP+ donor cells; (**C)** Leukocyte counts; (**D)** Erythrocyte counts; **(E)** Hemoglobin. Data for GFP+ donor cells, Leukocytes and Hemoglobin are shown as median ± IQR, data for Erythrocytes and Platelets are shown as mean ± SEM, unpaired T-tests * p<0.05, *** p<0.001 (n=11-27). Representative PB parameters of secondary recipients of *MUT-Hoxa1* BM that developed sAML: (**F)** total leukocytes excluding nucleated erythrocytes; (**G)** Erythrocytes; (**H)** Platelets; (**I)** Hemoglobin; (**J)** Mean corpuscular volume (MCV), n= 5. Representative PB smears and BM trephine H&E sections of a control mouse (**K** and **N)**, sAML (M1 subtype) in a recipient of *MUT-Hoxa1* BM (**L** and **O**), sAML (M6 subtype) in a recipient mouse of a separate *MUT-Hoxa1* BM cohort (**M** and **P**). Brightness and contrast were adjusted equally for visualization purposes. White arrows indicate AML blasts (**L, M**). Green arrows indicate nucleated red blood cells with associated dyserythropoietic changes (**M**). BM sections revealed an accumulation of blasts in sAML M1 subtype (**O**) and numerous nucleated red blood cells in sAML M6 subtype (**P**). Scale bar, 20 µm.

The majority of *MUT-Hoxa1* recipients developed sAML (9/10 separate cohorts, n= 4 per group). *MUT-Hoxa1* mice with sAML progression had a median overall survival of 10 months post-BMT (Figure 3A). These mice displayed variable leukocytosis, severe anemia and thrombocytopenia (Figure 3F-J) and had a normal karyotype. PB smears and BM sections confirmed sAML compared to MXIE control recipients (Figure 3K-P). Leukemic infiltration into non-hematopoietic tissues was also observed (supplemental Figure S1A-I). These were classified as AML without maturation (FAB AML-M1; Figure 3L and O) or erythroleukemia (FAB AML-M6; Figure 3M and P) in separate cohorts of mice.

Retroviral vector integration site analysis revealed that the majority of sAML occurring in *MUT-Hoxa1* recipients were likely to be independent of known co-operative integration site events, with the exception of a rapid sAML in one cohort due to a co-operating retroviral integration site at the *Meis1* gene (Table S2).

### DNA damage sensing and repair are downregulated in *WT-Hoxa1* and *MUT-Hoxa1* overexpressing MEPs and CMPs

Gene expression profiling was performed on MEPs and CMPs isolated from *WT-Hoxa1* and *MUT-Hoxa1* primary recipient mice at 16 weeks post-BMT. For MEPs, 79 genes were differentially expressed in *WT-Hoxa1* compared to MXIE control [expression log_2_ >1 with false discovery rate (q) <0.05], while 648 genes were altered in *MUT-Hoxa1* (Supplemental Dataset 1). For CMPs, 101 genes were differentially expressed in *WT-Hoxa1* and 223 genes differentially expressed in *MUT-Hoxa1* compared to MXIE (Supplemental Dataset 1). No genes were differentially expressed between *WT-Hoxa1* and *MUT-Hoxa1* MEPs or CMPs, however, *MUT-Hoxa1* demonstrated higher mean-fold changes than *WT-Hoxa1* for the majority of genes that were commonly dysregulated (Supplemental Dataset 1, Supplemental Table S3).

Quantitative Set Analysis of Gene Expression (QuSAGE) ^36^ analysis revealed that pathways affecting DNA replication and mismatch repair pathways were significantly downregulated in the *MUT-Hoxa1* MEPs and, to a lesser extent, *WT-Hoxa1* MEPs (Figure 4A-B, Supplemental Dataset 2). These pathways were also significantly downregulated in the *MUT-Hoxa1* CMPs (Figure 4C-D, Supplemental Dataset 3). We therefore assessed whether *MUT-Hoxa1*-overexpressing cells were more susceptible to DNA damage.

**Figure 4.**
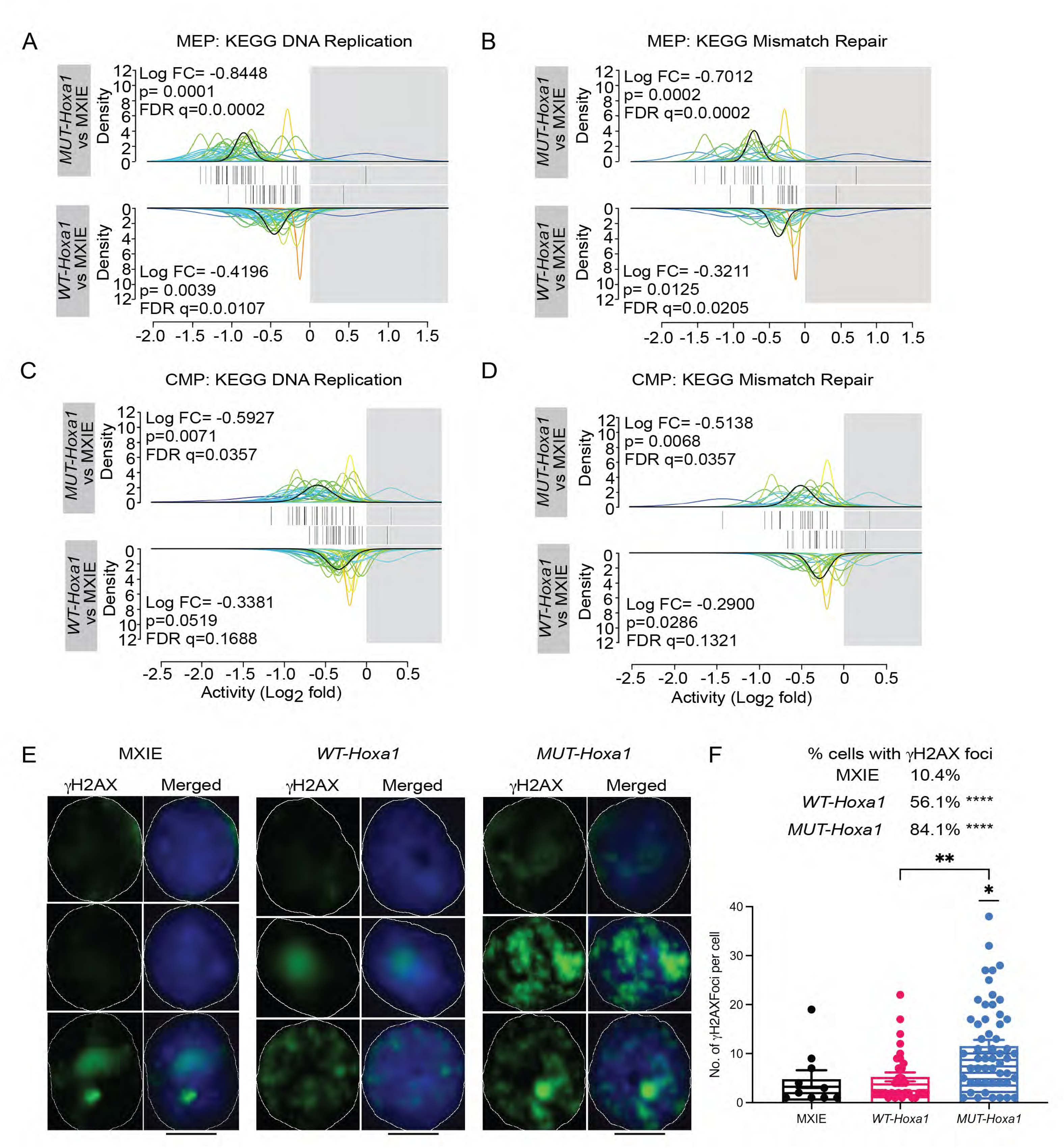
Altered DNA repair-related pathways in *WT-Hoxa1* and *MUT-Hoxa1* myeloid progenitors. QuSAGE analysis of 2 representative pathways enriched in *MUT-Hoxa1* and *WT-Hoxa1* myeloid progenitors compared to MXIE: (**A)** KEGG DNA replication in MEPs and (**B)** KEGG mismatch repair in MEPs. (**C)** KEGG DNA replication in CMPs and (**D)** KEGG mismatch repair in CMPs. The black curve line represents the average alteration for an individual pathway. Barcodes indicate the mean expression levels of individual genes in the set. Log FC= log-2-fold change, FDR= false discovery rate corrected p value (q). (**E)**, Representative γH2AX staining and (**F)**, percentage of cells with foci and quantitation of the number of γH2AX foci per cell in GFP+ MXIE, *WT-Hoxa1* and *MUT-Hoxa1* overexpressing cells cultured in cytokine-containing media for 2 weeks. Cells were imaged on a Nikon A1R laser scanning confocal microscope. Scale bar, 5 μm. Images have been thresholded, brightness and contrast were adjusted equally for visualization purposes, and all analyses were performed on raw data. Data in F are shown as median with interquartile range; *p<0.05, **p<0.01, ****p<0.0001 vs MXIE control or between groups designated by the bars. The Chi-Square test was used for comparisons of the percentages of cells with γH2AX foci; the numbers of γH2AX foci per cell were analyzed by one-way ANOVA with multiple comparisons. Data are pooled from two independent experiments.

Both the *WT-Hoxa1* and *MUT-Hoxa1*-overexpressing hematopoietic cells had a higher percentage of cells that contained γH2AX foci compared to MXIE (Figure 4E-F). *MUT-Hoxa1*-overexpressing cells had significantly more γH2AX foci per cell compared to both *WT-Hoxa1*-overexpressing cells and MXIE controls (Figure 4E-F). In contrast, no differences were observed in the numbers of γH2AX foci per cell in *WT-Hoxa1*-overexpressing cells compared to MXIE (Figure 4E-F).

### *HOXA1-FL* mRNA expression is upregulated in human MDS

Our data suggested a novel role for HOXA1 in MDS. *HOXA1* mRNA expression was therefore evaluated by *in silico* analysis of a microarray database of human MDS CD34^+^ cells (17 healthy controls, 183 MDS patients) ^32^. *HOXA1* mRNA expression was significantly increased in approximately 50% of MDS samples in comparison to healthy controls (Figure 5A). Amongst the 11 *HOXA* cluster genes, *HOXA1* displayed the highest fold-change in expression in MDS patient cells (Table S4). Furthermore, *HOXA1* was significantly upregulated in patients with MDS independent of chromosomal abnormalities (Figure 5B).

**Figure 5.**
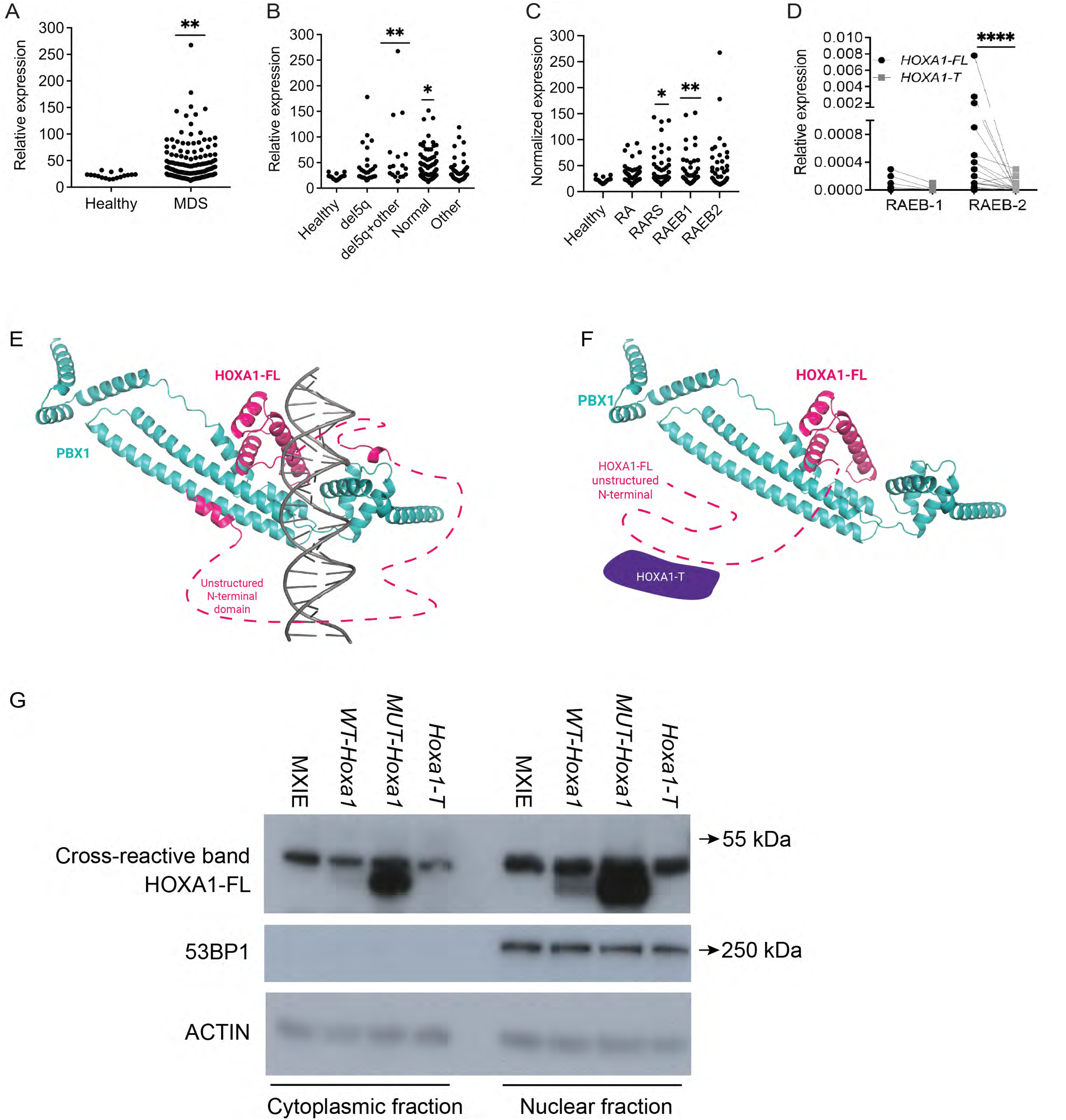
*HOXA1* is upregulated in CD34^+^ cells from MDS patients. Shown are (**A**) Expression of *HOXA1* transcripts in CD34^+^ BM cells from healthy and MDS patients, analyzed from ^32^ (182 MDS patients, 17 healthy controls). (**B)** Expression of *HOXA1* transcripts in CD34+ BM cells from patients with deletion 5q [del(5q) (n=29), del 5q + other cytogenetic abnormalities (n=18)], normal karyotype (n=17), and other abnormalities including trisomy 8 or monosomy 7 (n=42). (**C)** Expression of *HOXA1* transcripts in CD34^+^ BM cells for MDS subsets compared to control. (**D)** Relative expression of *HOXA1-FL* and *HOXA1-T* transcripts in CD34^+^ BM cells analyzed by Q-PCR. Lines connect the expression of *HOXA1-FL* and *HOXA1-T* in individual samples (10 RAEB-1 MDS patients, 5 of which expressed HOXA1 transcripts and 25 RAEB-2 MDS patients, 20 of which expressed HOXA1 transcripts). *p < 0.05; **p < 0.01, ***p < 0.001 vs. healthy controls. The Mann-Whitney two-tailed U test was used for statistical analysis of data in (**A)**, one-way ANOVA with multiple comparisons was used for statistical analysis of data in (**B)** and (**C)**, data comparing (**D)**, *HOXA1-FL* vs *HOXA1-T* within a patient sample were analyzed using paired two-tailed Wilcoxon signed rank test; data comparing *HOXA1-FL* (p=0.055) or *HOXA1-T* (p=0.68) in RAEB-1 versus RAEB-2 patients were analyzed using the Mann-Whitney two-tailed U test. The proposed 3D atomic structure of transcriptionally active HOXA1 is shown in ribbon representation in (**E);** HOXA1-FL and HOXA1-T are proposed to form heterodimers via the N-terminus domain shown in (**F)**. Represented in grey: double-stranded DNA, magenta: HOXA1-FL, teal: PBX1 co-factor, and purple: HOXA1-T. (**G**) Representative Western blot showing HOXA1-FL expression in cytoplasmic and nuclear fractions of GP+E-86 ecotropic packaging cells overexpressing the indicated vector. The same Western blot was stripped and reprobed for nuclear control (53BP1) and loading control (ACTIN). Brightness and contrast have been adjusted equally for visualization purposes.

*HOXA1* expression was significantly increased in two MDS subtypes assessed on the microarray, classified according to the French-American-British (FAB) system: RA with ringed sideroblasts (RARS) and RA with excess blasts 1 (RAEB-1) (Figure 5C). We assessed the mRNA expression of the two *HOXA1* isoforms in BM CD34^+^ cells isolated from a cohort of high-risk (RAEB-1 and RAEB2) MDS patients^37^ (Table S5). 20 of the 25 RAEB-2 patient samples analyzed expressed *HOXA1-FL* and 13 of the 20 patients did not express *HOXA1-T*, with significantly higher expression of *HOXA1-FL* observed (Figure 5D). Analyses of RNA-seq data of an independent cohort of MDS patients (23 healthy, 44 MDS) ^33^ revealed that *HOXA1-FL* was significantly upregulated compared to *HOXA1-T* in patients with MDS classified as intermediate 1 (Int 1) risk using the international prognosis scoring system (IPSS) for MDS (supplemental Figure S3A).

Somatic mutation data were available from the published RNA-seq series^33^ and the RAEB- 1 and RAEB-2 patients shown in Figure 5D^37^, therefore the mutation patterns were assessed together with *HOXA1-FL* and *HOXA1-T* expression (supplemental Figure S3B-C). There were no differences in the distribution of somatic mutations within the different functional categories based on *HOXA1-FL* expression (supplemental Figure S3B-C).

The 3D atomic structure of HOXA1 has not been solved, however, it shares 78% homology with HOXB1, for which the structure has been determined^38^. We threaded amino acid residues 202-290 of the HOXA1 sequence onto the solved structure of HOXB1^38^ in complex with DNA and its co-factor PBX1, then minimized the complex utilizing molecular dynamics. An independent AlphaFold model ^39^ showed the same structural fold. The remaining 201 amino acids of the N-terminus of HOXA1-FL were predicted to be unstructured (Figure 5E). HOXA1-FL and HOXA1-T have been proposed to form heterodimers via the unstructured N-terminal domain ^6^. Our modeling suggested that this would cause a structural rearrangement, such that DNA can no longer bind to the complex (Figure 5F), implying that HOXA1-T is a negative regulator of transcriptional activation by HOXA1-FL. In contrast, the lack of HOXA1-T, as occurs in the *MUT-Hoxa1* model, would likely result in unimpeded binding of HOXA1-FL and its co-factor (PBX1) to DNA, enhancing the transcriptional activity of HOXA1-FL. Indeed, western blotting of protein isolated from the overexpressing cell lines revealed that HOXA1-FL was upregulated in both the nucleus and the cytoplasm of *MUT-Hoxa1* cells compared to *WT-Hoxa1*, *Hoxa1-T* and MXIE cells (Figure 5G). HOXA1-FL was also noticeably increased in the cytoplasm and nucleus of the *WT-Hoxa1* cells compared to MXIE and *Hoxa1-T* (Figure 5G).

### Conditional knock-in mouse models of *WT-Hoxa1* and *MUT-Hoxa1* develop cytopenias

Having confirmed a potential role for HOXA1 in MDS pathogenesis, conditional *Rosa26*-lox-stop-lox *WT-Hoxa1* and *MUT-Hoxa1* knock-in mice were generated (Figure 6A-B). The expression of *Hoxa1-FL* and *Hoxa1-T* was confirmed in sorted hematopoietic cells (supplemental Figure S4A-C). *WT-Hoxa1* and *MUT-Hoxa1* knock-ins were crossed to tamoxifen-inducible hematopoietic specific *hScl-*Cre^ERT^ mice ^30^. To determine if the amount of *Hoxa1-FL* overexpressed in the hematopoietic cells determined the phenotype, we generated heterozygous (*WT-Hoxa1^ki/+^*and *MUT-Hoxa1^ki/+^*) and homozygous (*WT-Hoxa1^ki/ki^*and *MUT-Hoxa1^ki/ki^*) knock-ins. 4-week-old mice were fed tamoxifen chow to induce the knock-ins.

**Figure 6.**
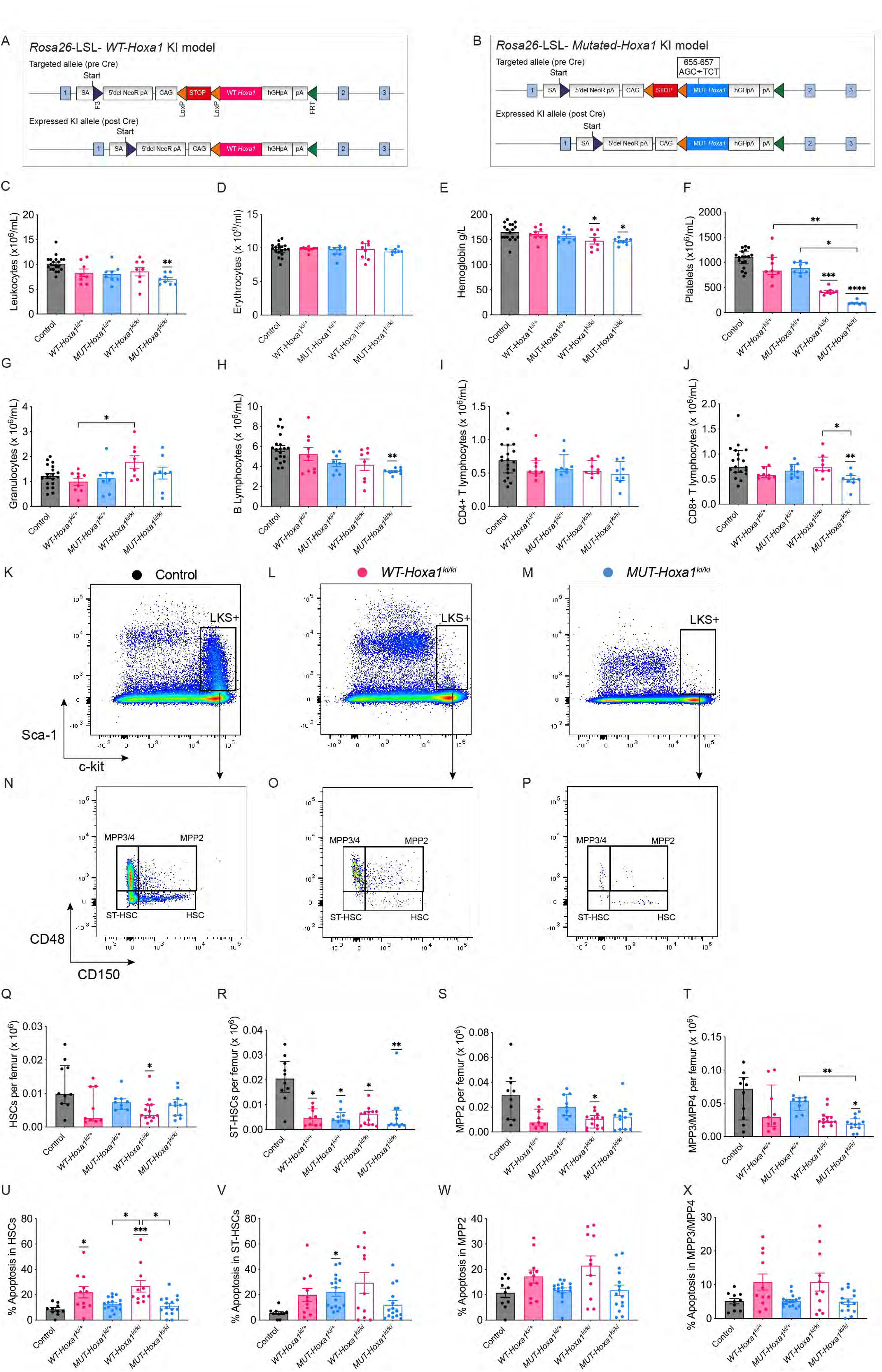
Conditional *Hoxa1* knock-in mice developed thrombocytopenia accompanied by reduced HSCs and MPPs. Schematic diagram of the gene targeting strategy for conditional knock-in mice of: (**A)** *WT-Hoxa1* and (**B)** *MUT-Hoxa1* expressed at the *R26* locus. PB counts of *WT-Hoxa1^ki/+^, MUT-Hoxa1^ki/+^, WT-Hoxa1^ki/ki^, MUT-Hoxa1^ki/ki^* and Cre^+^ controls at 4 months after induction of knock-in: (**C)** Leukocytes, (**D**) Erythrocytes, (**E**) Hemoglobin and (**F)** Platelets. Numbers of PB leukocyte subsets: (**G)** granulocytes, (**H)** B lymphocytes, (**I)** CD4^+^ T lymphocytes and (**J)** CD8^+^ T lymphocytes (n>6). Representative flow cytometry plots of Cre^+^ control, *WT-Hoxa1^ki/ki^* and *MUT-Hoxa1^ki/ki^* BM showing: (**K-M**): LKS+ profiles and (**N-P**): HSPC profiles within LKS+ cells. Numbers of: (**Q)** HSCs, (**R)** ST-HSCs, (**S)** MPP2, (**T)** MPP3/MPP4 in BM (n=9-13). Percentages of apoptotic cells in: **(U)** HSCs, (**V)** ST-HSCs, (**W)** MPP2, (**X)**, MPP3/4 (n=10-16). All data are shown as median ± IQR except C, E, G, H and X which are mean ± SEM; *p<0.05; **p<0.01; ***p<0.001; ****p<0.0001 vs control, analyzed by one-way ANOVA with multiple comparisons. Data are pooled from multiple experiments.

Mice were analyzed at 4 months post-tamoxifen initiation, and changes in hematopoiesis were compared to tamoxifen-treated *hScl-Cre*^ERT^ Cre^+^ *Rosa26* wildtype control mice. *MUT-Hoxa1^ki/ki^* mice had significantly lower PB leukocyte counts (Figure 6C). PB erythrocyte numbers were unchanged, however, both *WT-Hoxa1^ki/ki^* and *MUT-Hoxa1^ki/ki^* mice had significantly reduced hemoglobin (Figure 6D-E). Furthermore *WT-Hoxa1^ki/ki^* and *MUT-Hoxa1^ki/ki^* mice developed profound thrombocytopenia, and *MUT-Hoxa1^ki/ki^* mice had significantly fewer platelets compared to both the *WT-Hoxa1^ki/+^*and *MUT-Hoxa1^ki/+^* mice (Figure 6F).

There were significantly increased PB granulocytes in *WT-Hoxa1^ki/ki^* mice (Figure 6G) and significant reductions in the numbers of PB B lymphocytes in the *MUT-Hoxa1^ki/ki^*mice (Figure 6H). The numbers of PB CD4^+^ T cells were unaltered (Figure 6I), however, the numbers of CD8^+^ T cells were significantly reduced in *MUT-Hoxa1^ki/ki^*mice compared to both control and *WT-Hoxa1^ki/ki^* mice (Figure 6J).

There were striking reductions of HSPCs in all *Hoxa1* genotypes compared to Cre+ controls (Figure 6K-T). The numbers of SLAM HSCs ^40,41^ were significantly reduced in *WT-Hoxa1^ki/ki^* mice (Figure 6Q) and the numbers of ST-HSCs were significantly reduced in all *Hoxa1* knock-in genotypes compared to Cre^+^ controls (Figure 6R). The numbers of MPP2 were significantly reduced in *WT-Hoxa1^ki/ki^* mice (Figure 6S). Significantly reduced numbers of MPP3/MPP4 were observed in *MUT-Hoxa1^ki/ki^* mice compared to both control and *MUT-Hoxa1^ki/+^* mice (Figure 6T).

There were significantly increased percentages of apoptotic cells in *WT-Hoxa1^ki/+^* and *WT-Hoxa1^ki/ki^* SLAM HSCs compared to all other genotypes (Figure 6U). There were also significantly increased proportions of apoptotic cells in *MUT-Hoxa1^ki/+^* ST-HSCs compared to Cre+ controls (Figure 6V), but no significant differences in apoptosis in the MPP populations (Figure 6W-X).

Somatic mutations were observed in total BM cells obtained from all mice, including healthy Cre^+^ controls, consistent with recent findings^42^. There were no differences in mutation burden in *Hoxa1* mice compared to Cre^+^ controls (supplemental Figure S4D-E). All mice had mutations associated with clonal hematopoiesis^43^, including *Sf3b1* and *Ezh2* (supplemental Figure S4D).

### Wildtype recipients of *WT-Hoxa1* and *MUT-Hoxa1* knock-in bone marrow develop MDS

BM cells from Cre^+^ controls, *WT-Hoxa1^ki/+^*, *MUT-Hoxa1^ki/+^, WT-Hoxa1^ki/ki^* and *MUT-Hoxa1^ki/ki^*mice were transplanted into wildtype recipients and monitored for MDS progression (Figure 7A-E). MDS was observed in *Hoxa1* recipients commencing at 12 months post-BMT (Figure 7A). Significant trilineage dysplasia was demonstrated on morphological assessment of peripheral blood smears and BM of mice from all knock-in genotypes (Figure 7C-E). This included hypersegmented PB neutrophils (Figure 7C), marked erythroid dysplasia (Figure 7D), and significant megakaryocytic atypia (Figure 7E). There were no increases in blasts. The *Hoxa1* knock-in mice therefore fulfill the diagnostic criteria for MDS, with features reminiscent of myelodysplasia with multilineage dysplasia (MDS-MLD).

**Figure 7.**
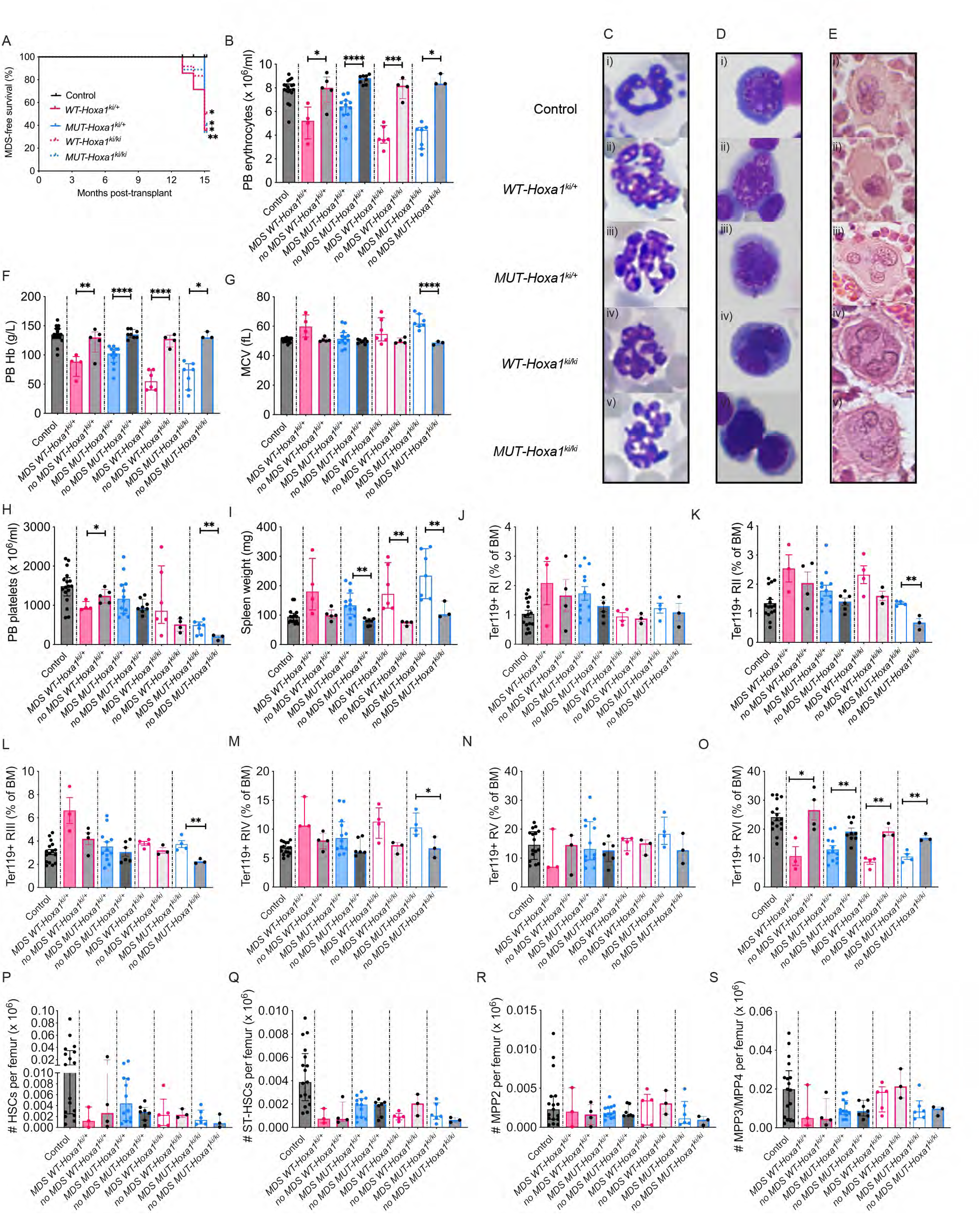
Recipients of *WT-Hoxa1* and *MUT-Hoxa1* knock-in BM develop MDS. Shown are: (**A)** Kaplan-Meier curve of MDS-free survival in transplant recipients of *WT-Hoxa1^ki/+^, MUT-Hoxa1^ki/+^, WT-Hoxa1^ki/ki^, MUT-Hoxa1^ki/ki^* and Cre^+^ control donor BM. All data are shown from transplant recipients analyzed at 12-15 months post-BMT. (**B**) Peripheral blood erythrocytes. (**C-E**): Representative PB smears, BM cytospins and BM trephine morphology of trilineage hematopoiesis in transplant recipients of (i) Cre^+^ control, (ii) *WT-Hoxa1^ki/+^,* (iii) *MUT-Hoxa1^ki/+^,* (iv) *WT-Hoxa1^ki/ki^ and* (v) *MUT-Hoxa1^ki/ki^* donor BM are shown in: (**C**), granulocytic lineage: (i) normal neutrophils (ii – v) dysplastic hypersegmented neutrophils observed in recipients of knock-in BM; (**D**), erythroid lineage: (i) normal erythroblast, dyserythropoietic changes observed in recipients of knock-in BM (ii-v) included binucleation, nuclear budding, internuclear bridging and multinucleation and (**E**), megakaryocytic lineage: (i) normal megakaryocyte (ii – v) dysplastic megakaryocytes with dispersed nuclei and hypolobated forms were seen in recipients of knock-in BM. Original magnification x 400. PB parameters: (**F**) Hemoglobin, (**G**) Mean Corpuscular Volume (MCV), (**H**) Platelet counts. (**I**) spleen weights. Shown are percentages of erythrocyte subsets in BM: (**J**), RI proerythroblasts, (**K**) RII basophilic erythroblasts, (**L**) RIII polychromatic erythroblasts, (**M**) RIV orthochromatic erythroblasts, (**N**) RV reticulocytes, (**O**) RVI mature erythrocytes. Shown are the numbers of: (**P**) HSC, (**Q**), ST-HSC, (**R)**, MPP2 and (**S**) MPP3/MPP4 in BM. Kaplan-Meier curve was analyzed with log-rank (Mantel–Cox) test, all other data were analyzed by unpaired T-tests between mice with MDS and mice without MDS within a given genotype, Cre^+^ Control data are shown for visual comparisons. Data in B, F, I, N, M, P and R are shown as median ± IQR, data in G, H, J, K, L, O and S are shown as mean ± SEM; *p<0.05; **p<0.01; ****p<0.0001 between the MDS and no MDS cohorts. Data are pooled from four separate experiments, n=3-17.

At 15 months post-transplant all remaining cohorts were euthanized for analysis. This enabled a comparison of phenotypes in mice that developed MDS versus mice that did not, revealing phenotypes that were MDS-specific versus those that had developed due to the effects of *Hoxa1* on hematopoiesis.

Within each genotype, all mice that developed MDS had significantly reduced PB erythrocyte counts (Figure 7B) and hemoglobin (Figure 7F) compared to mice that did not develop MDS. Recipients of *MUT-Hoxa1^ki/ki^* MDS had macrocytic anemia (Figure 7G). PB platelet counts were low in all cohorts transplanted with *Hoxa1* BM (Figure 7H). Mice with *WT-Hoxa1^ki/+^* MDS had PB neutropenia whereas mice with *MUT-Hoxa1^ki/ki^*MDS developed PB B lymphopenia compared to recipients that did not develop MDS (supplemental Figure S5A-E). Recipients of *MUT-Hoxa1^ki/ki^* BM that did not develop MDS had significantly increased numbers of PB granulocytes (supplemental Figure S5B).

There were no differences in the BM leukocyte cellularities, granulopoiesis or CD4^+^ T lymphocytes in the BM (supplemental Figure S5F-I). MDS recipients of *WT-Hoxa1^ki/+^* BM had significantly reduced numbers of BM CD8^+^ T lymphocytes compared to those without MDS (supplemental Figure S5J).

Significant splenomegaly was observed in recipients of *MUT-Hoxa1^ki/+^*, *WT-Hoxa1^ki/ki^ and MUT-Hoxa1^ki/ki^* BM that developed MDS (Figure 7I). This was accompanied by ineffective erythropoiesis in the BM of *Hoxa1* recipients that developed MDS (Figure 7J-O). There were significantly increased fractions of RII-RIV immature erythroid cells^34^ in *MUT-Hoxa1^ki/ki^* MDS recipients compared to those without MDS (Figure 7K-M). All MDS recipients had significantly reduced proportions of RVI mature erythrocytes (Figure 7O).

The numbers of HSPCs were reduced in *Hoxa1* recipients, however, there were no differences observed in HSPC content between mice that had developed MDS versus those that did not (Figure 7P-S).

Whole exome sequencing of BM from the mice revealed that, in addition to mutations observed in pre-MDS donors (supplemental Figure S4D), other mutations, including *Dnmt3a*^43^, were observed in significant cohorts of mice, including controls (supplemental Figure S5L-M). There was no specific mutational signature in *Hoxa1* knock-in recipients that developed MDS compared to either controls or *Hoxa1* recipients that did not develop MDS (supplemental Figure S5K-M), consistent with that observed in MDS patient samples (supplemental Figure S3).

## Discussion

Increased expression of the transcriptionally active full length HOXA1 in mice perturbed hematopoiesis and caused MDS. Of clinical significance, increased expression of *HOXA1* was observed in CD34^+^ BM cells from between 30-50% of MDS patients from two independent cohorts. The main change in *HOXA1* expression observed in MDS patients was increased expression of the homeobox-containing *HOXA1-FL* isoform. However, we also found evidence of loss of expression of the homeoboxless *HOXA1-T* in MDS patients, in particular in patients with RAEB-2 and Int 1 risk disease by IPSS. These data suggest that *HOXA1* is frequently dysregulated in MDS, consistent with previous suggestions that there are common mechanisms that contribute to MDS ^44^.

Collectively, our data suggest that HOXA1-T is a negative regulator of HOXA1-FL. Overexpression of the truncated *Hoxa1-T* in hematopoietic cells had opposing effects on repopulation of ST-HSCs and MPPs *in vivo* compared to *WT-Hoxa1* and *MUT-Hoxa1* (Figure 2O-P). Computational modelling suggested that HOXA1-T binds to HOXA1-FL and changes the conformation, regulating the ability of HOXA1-FL to activate gene transcription. Consistent with this, we observed more HOXA1-FL protein in both the nucleus and cytoplasm of a *MUT-Hoxa1* compared to *WT-Hoxa1* overexpressing cell line (Figure 5). Furthermore, there were no differentially expressed transcripts between *MUT-Hoxa1* compared to *WT-Hoxa1* MEPs or CMPs, however, for the majority of the transcripts that were commonly dysregulated, *MUT-Hoxa1* progenitors had higher mean-fold changes compared to *WT-Hoxa1* progenitors (Supplemental Datasets 2 and 3).

The retroviral and knock-in *Hoxa1* mice provide reproducible models of MDS that differ in their phenotypic severity and propensity to develop sAML. The phenotypes observed in the *WT-Hoxa1* models were reminiscent of lower risk MDS, having cytopenias with associated dysplasia, accompanied by increased apoptosis observed in the HSPCs. In contrast, *MUT-Hoxa1* recipients resembled higher risk MDS with pancytopenia, increased DNA damage and progression to sAML, the latter of which was observed in the retroviral model.

In summary, we have established novel, faithful mouse models of MDS based on dysregulated *Hoxa1* isoform expression. Our mouse models cover a spectrum of disease progression, including pre-MDS, specifically CHIP (single knock-ins) and CCUS (double knock-ins). The ability to compare phenotypes of mice that that develop MDS versus mice that do not within the same time period will enable the distinction of key factors that cause MDS. With alterations in *HOXA1* expression observed in up to 50% of MDS patients, these mouse models will be valuable pre-clinical tools to further understand MDS pathogenesis, as well as facilitating the identification of potential novel therapies for this malignant disease.

## Acknowledgements

We thank Jean Hendy, Stewart Fabb, Ana Maluenda, Emma Baker, Julie Quach, Tanja Jovic and Mannu Walia for excellent technical assistance, Jörg Heierhorst and Harshal Nandurkar for excellent discussions, the SVH Bioresources Centre for care of experimental animals and SVI FACS Facility for FACS sorting. This work was supported in part by grants from the Leukaemia Foundation (to LEP, MW and MWP), the Cancer Council Victoria (to LEP, MW and CRW, GNT1104715), Cancer Australia (to LEP and JKH, GNT2010541), National Health and Medical Research Council of Australia (to LEP, GNT2028698), the Zig Inge Foundation (to LEP), the Leukaemia Foundation (to MW) and the Victorian State Government Operational Infrastructure Support Program (to St. Vincent’s Institute). ST was the recipient of a Leukaemia Foundation Clinical PhD Scholarship supported by Andrew Cadigan in honour of Chris Simpson. CSG was the recipient of an Australian Government Research Training Program PhD Scholarship. JQT was supported by a Cancer Council Victoria Postdoctoral Research Fellowship. JKH was supported by a Postdoctoral Fellowship from the Leukaemia Foundation in partnership with the Cure Cancer Australian Foundation. CRW was supported by the Philip Desbrow Senior Research Fellowship of the Leukaemia Foundation and a Victorian Cancer Agency Fellowship. MW was supported by a Victorian Cancer Agency Clinical Research Fellowship. LEP was a Senior Research Fellow (GNT1003339) and MWP was a Senior Principal Research Fellow of the National Health and Medical Research Council of Australia (GNT1117183). Dedicated to the memory of Andrew Cadigan and Chris Simpson.

## Author Contributions

LEP conceived the study. LEP, ST and CJ designed and performed experiments, analyzed data and co-wrote the manuscript; MLS, AMC, SCL, KS, GT, SG, SM, CSG, JQT, MFS, KLR and JKH performed experiments and analyzed data; AU, AGK, GJM, MT, EH-L, JEP and LJG provided valuable reagents, MWP, DMcC and CRW analyzed data, MW designed experiments and analyzed data, All authors read and approved the final manuscript.

## Data Availability Statement

Raw data microarray files are available at GEO using accession number GSE62853 for MEPs and GSE227775 for CMPs. All other data that support the findings of this study are available from the corresponding author (LEP).

## References

1. Shen WF, Montgomery JC, Rozenfeld S, et al. AbdB-like Hox proteins stabilize DNA binding by the Meis1 homeodomain proteins. Mol Cell Biol. 1997;17(11):6448–6458.

2. Shen WF, Rozenfeld S, Kwong A, Kom ves LG, Lawrence HJ, Largman C. HOXA9 forms triple complexes with PBX2 and MEIS1 in myeloid cells. Mol Cell Biol. 1999;19(4):3051–3061.

3. Mann RS. The specificity of homeotic gene function. Bioessays. 1995;17(10):855–863.

4. Komuves LG, Shen WF, Kwong A, et al. Changes in HOXB6 homeodomain protein structure and localization during human epidermal development and differentiation. Dev Dyn. 2000;218(4):636–647.

5. LaRosa GJ, Gudas LJ. Early retinoic acid-induced F9 teratocarcinoma stem cell gene ERA-1: alternate splicing creates transcripts for a homeobox-containing protein and one lacking the homeobox. Mol Cell Biol. 1988;8(9):3906–3917.

6. Fernandez CC, Gudas LJ. The truncated Hoxa1 protein interacts with Hoxa1 and Pbx1 in stem cells. J Cell Biochem. 2009;106(3):427–443.

7. Fujimoto S, Araki K, Chisaka O, Araki M, Takagi K, Yamamura K. Analysis of the murine Hoxa-9 cDNA: an alternatively spliced transcript encodes a truncated protein lacking the homeodomain. Gene. 1998;209(1-2):77–85.

8. Hong YS, Kim SY, Bhattacharya A, Pratt DR, Hong WK, Tainsky MA. Structure and function of the HOX A1 human homeobox gene cDNA. Gene. 1995;159(2):209–214.

9. Stadler CR, Vegi N, Mulaw MA, et al. The leukemogenicity of Hoxa9 depends on alternative splicing. Leukemia. 2014;28:1838–1843.

10. Murphy P, Hill RE. Expression of the mouse labial-like homeobox-containing genes, Hox 2.9 and Hox 1.6, during segmentation of the hindbrain. Development. 1991;111(1):61–74.

11. Chisaka O, Musci TS, Capecchi MR. Developmental defects of the ear, cranial nerves and hindbrain resulting from targeted disruption of the mouse homeobox gene Hox-1.6. Nature. 1992;355(6360):516–520.

12. Lufkin T, Dierich A, LeMeur M, Mark M, Chambon P. Disruption of the Hox-1.6 homeobox gene results in defects in a region corresponding to its rostral domain of expression. Cell. 1991;66(6):1105–1119.

13. Sauvageau G, Lansdorp PM, Eaves CJ, et al. Differential expression of homeobox genes in functionally distinct CD34+ subpopulations of human bone marrow cells. Proc Natl Acad Sci U S A. 1994;91(25):12223–12227.

14. Lebert-Ghali CE, Fournier M, Dickson GJ, Thompson A, Sauvageau G, Bijl JJ. HoxA cluster is haploinsufficient for activity of hematopoietic stem and progenitor cells. Exp Hematol. 2010;38(11):1074–1086 e1071–1075.

15. Argiropoulos B, Humphries RK. Hox genes in hematopoiesis and leukemogenesis. Oncogene. 2007;26(47):6766–6776.

16. Borrow J, Shearman AM, Stanton VP, Jr., et al. The t(7;11)(p15;p15) translocation in acute myeloid leukaemia fuses the genes for nucleoporin NUP98 and class I homeoprotein HOXA9. Nat Genet. 1996;12(2):159–167.

17. Raza-Egilmez SZ, Jani-Sait SN, Grossi M, Higgins MJ, Shows TB, Aplan PD. NUP98-HOXD13 gene fusion in therapy-related acute myelogenous leukemia. Cancer Res. 1998;58(19):4269–4273.

18. Soulier J, Clappier E, Cayuela JM, et al. HOXA genes are included in genetic and biologic networks defining human acute T-cell leukemia (T-ALL). Blood. 2005;106(1):274–286.

19. Lebert-Ghali CE, Fournier M, Kettyle L, Thompson A, Sauvageau G, Bijl JJ. Hoxa cluster genes determine the proliferative activity of adult mouse hematopoietic stem and progenitor cells. Blood. 2016;127(1):87–90.

20. Arber DA, Orazi A, Hasserjian R, et al. The 2016 revision to the World Health Organization classification of myeloid neoplasms and acute leukemia. Blood. 2016;127(20):2391–2405.

21. Tan SY, Smeets MF, Chalk AM, et al. Insights into myelodysplastic syndromes from current preclinical models. World Journal of Hematology. 2016;5(1):1–22.

22. Li H, Hu F, Gale RP, Sekeres MA, Liang Y. Myelodysplastic syndromes. Nat Rev Dis Primers. 2022;8(1):74.

23. Saygin C, Godley LA. Genetics of Myelodysplastic Syndromes. Cancers (Basel). 2021;13(14).

24. Merz AMA, Platzbecker U. Beyond the horizon: emerging therapeutic approaches in myelodysplastic neoplasms. Exp Hematol. 2024;130:104130.

25. Mina A, Pavletic S, Aplan PD. The evolution of preclinical models for myelodysplastic neoplasms. Leukemia. 2024;38(4):683–691.

26. Valent P. ICUS, IDUS, CHIP and CCUS: Diagnostic Criteria, Separation from MDS and Clinical Implications. Pathobiology. 2019;86(1):30–38.

27. Osman A. When are idiopathic and clonal cytopenias of unknown significance (ICUS or CCUS)? Hematology Am Soc Hematol Educ Program. 2021;2021(1):399–404.

28. Robbins PB, Yu XJ, Skelton DM, et al. Increased probability of expression from modified retroviral vectors in embryonal stem cells and embryonal carcinoma cells. J Virol. 1997;71(12):9466–9474.

29. Purton LE, Dworkin S, Olsen GH, et al. RARgamma is critical for maintaining a balance between hematopoietic stem cell self-renewal and differentiation. J Exp Med. 2006;203(5):1283–1293.

30. Gothert JR, Gustin SE, Hall MA, et al. In vivo fate-tracing studies using the Scl stem cell enhancer: embryonic hematopoietic stem cells significantly contribute to adult hematopoiesis. Blood. 2005;105(7):2724–2732.

31. Green AC, Tjin G, Lee SC, et al. The characterization of distinct populations of murine skeletal cells that have different roles in B lymphopoiesis. Blood. 2021;138:304–317.

32. Pellagatti A, Cazzola M, Giagounidis A, et al. Deregulated gene expression pathways in myelodysplastic syndrome hematopoietic stem cells. Leukemia. 2010;24(4):756–764.

33. Im H, Rao V, Sridhar K, et al. Distinct transcriptomic and exomic abnormalities within myelodysplastic syndrome marrow cells. Leuk Lymphoma. 2018;59(12):2952–2962.

34. Liu J, Zhang J, Ginzburg Y, et al. Quantitative analysis of murine terminal erythroid differentiation in vivo: novel method to study normal and disordered erythropoiesis. Blood. 2013;121(8):e43–49.

35. Bach C, Buhl S, Mueller D, Garcia-Cuellar MP, Maethner E, Slany RK. Leukemogenic transformation by HOXA cluster genes. Blood. 2010;115(14):2910–2918.

36. Yaari G, Bolen CR, Thakar J, Kleinstein SH. Quantitative set analysis for gene expression: a method to quantify gene set differential expression including gene-gene correlations. Nucleic Acids Res. 2013;41(18):e170.

37. Unnikrishnan A, Papaemmanuil E, Beck D, et al. Integrative Genomics Identifies the Molecular Basis of Resistance to Azacitidine Therapy in Myelodysplastic Syndromes. Cell Rep. 2017;20(3):572–585.

38. Piper DE, Batchelor AH, Chang CP, Cleary ML, Wolberger C. Structure of a HoxB1-Pbx1 heterodimer bound to DNA: role of the hexapeptide and a fourth homeodomain helix in complex formation. Cell. 1999;96(4):587–597.

39. Jumper J, Evans R, Pritzel A, et al. Highly accurate protein structure prediction with AlphaFold. Nature. 2021;596(7873):583–589.

40. Oguro H, Ding L, Morrison SJ. SLAM family markers resolve functionally distinct subpopulations of hematopoietic stem cells and multipotent progenitors. Cell Stem Cell. 2013;13(1):102–116.

41. Purton LE. Adult murine hematopoietic stem cells and progenitors: an update on their identities, functions, and assays. Exp Hematol. 2022;116:1–14.

42. Kapadia CD, Williams N, Dawson KJ, et al. Clonal dynamics and somatic evolution of haematopoiesis in mouse. Nature. 2025;641(8063):681–689.

43. Dunn WG, McLoughlin MA, Vassiliou GS. Clonal hematopoiesis and hematological malignancy. J Clin Invest. 2024;134(19).

44. Pellagatti A, Armstrong RN, Steeples V, et al. Impact of spliceosome mutations on RNA splicing in myelodysplasia: dysregulated genes/pathways and clinical associations. Blood. 2018;132(12):1225–1240.

