## Supplemental Data for "Dysregulated expression of *Hoxa1* isoforms in hematopoietic stem and progenitor cells causes myelodysplastic syndromes"

#### Supplemental Information

#### Supplemental Materials and Methods

##### RNA extraction and QPCR

Total RNA extraction and DNase treatment was carried out using the RNeasy Micro kit (Qiagen) or ISOLATE II RNA Micro kit (Bioline). cDNA was prepared using the AffinityScript QPCR cDNA synthesis kit with random primers (Agilent Technologies). QPCR was performed using the SensiMix II Probe Kit (Bioline) or Brilliant II Sybr Green QPCR master mix (Agilent Technologies) on the MX3000P Multiplex Quantitative PCR system (Stratagene). Primer/probe sequences used for QPCR are in Supplemental Table S6. The relative gene expression was calculated by the $2^{-\Delta CT}$ method using beta-2-microglobulin as reference gene.

**Transduction Efficiencies**

All transduction efficiencies were measured immediately post-transduction by GFP expression of the transduced cells. The average transduction efficiencies (%GFP in the cultures) of each of the BM cells were as follows (mean ± SEM, representative from n=3-4 separate transductions): MXIE: 29.22 ± 2.62; *WT-Hoxa1*: 7.72 ± 0.55; *MUT-Hoxa1*: 7.89 ± 0.80; *Hoxa1-T*: 10.38 ± 1.74. MXIE-transduced cells had significantly increased transduction efficiencies compared to all other groups ($p<0.0001$).

***In vitro* cultures and Colony-Forming Cell (CFC) Assay**

Cell cultures were initiated with 10,000 GFP+ cells (sorted post-transduction) per population cultured in triplicate as previously described [1]. At weekly intervals, cell counts were assessed using trypan blue and 10,000 cells were replated. CFCs on 100 $LKS^+$ GFP+ cells (sorted post-transduction) were performed as previously described [2].

**Retroviral Integration Analysis**

Retroviral integration site was determined by splinkerette PCR method [3]. 1-5µg of genomic DNA was digested using Sau3AI enzyme (New England Biolabs). Digested DNA was ligated to annealed splinkerette oligos with T4 DNA ligase (New England Biolabs). Ligated DNA was digested using Eco RV enzyme (New England Biolabs) to remove any splinkerette oligos ligated onto the LTR end of the Sau3AI digested genomic DNA to prevent amplification of internal retroviral fragments. Following DNA clean up, primary and secondary (nested) PCR reactions were carried out. Sequencing of the PCR products was carried out using standard conditions. The PCR primers used are listed in Supplemental Table S6.

**Analyses of MEP and CMP microarray data.**

MEP and CMP data were analysed separately. Raw CEL files were normalised with RMA [4] in R [v3.4] using oligo [v1.40.2, core probesets] [5] and annotated with a genepattern chip annotation file (Affymetrix_MoGene_1_1_st_v1.na33.2.mm9.transcript.chip). One each MEP and CMP *WT-Hoxa1* sample (r3) was discarded, as it did not cluster with the other samples from the *WT-Hoxa1* group and was subsequently shown to be pre-leukemic (data not shown).

For each comparison (*WT-Hoxa1* vs. MXIE, *MUT-Hoxa1* vs. MXIE, *MUT-Hoxa1* vs *WT-Hoxa1*), only those probesets detected above background (greater than the median of the control probes on each chip + 1 standard deviation) in at least 75% of a sample class were used (> 6,000 probesets each). Differentially expressed genes were calculated by Limma [6] with array weighting. The threshold for significant differential expression was FDR < 0.05 and abs(logFC) > 1. All data are provided in Supplemental Dataset 1.

Pathway analysis was performed using Ingenuity Pathway Analysis software (Qiagen, Redwood, CA). QuSAGE was used to identify differential genesets [7]. For each comparison (*WT-Hoxa1* vs MXIE, *MUT-Hoxa1* vs MXIE), gene expression data was collapsed to gene level using the most informative probe and run against the hallmark and c2.all gene sets using QuSAGE (n.points = 2^16, var. method = Welch's, var.equal = no). The sets from the MSigDB (version 5.2, Liberzon et al. 2011) were first processed into mouse orthologues as calculated by WEHI (http://bioinf.wehi.edu.au/software/MSigDB/). Briefly, the current MSigDB v5.2 xml file was downloaded. Human Entrez Gene IDs were mapped to Mouse Entrez Gene IDs, using the HGNC Comparison of Orthology Predictions (HCOP) (downloaded 11 October 2016). EntrezIDs were subsequently converted to symbols for visualization.

#### γH2AX immunofluorescence studies

Transduced MXIE, *WT-Hoxa1* and *MUT-Hoxa1* BM cells were sorted for GFP then cultured for 2 weeks. Cells were fixed in 2% paraformaldehyde prior to performing cytospins and staining for γH2AX as previously described [8,9]. Cells were stained with primary rabbit anti-mouse γH2AX (Cell Signaling Technology), secondary biotinylated donkey anti-rabbit (Jackson ImmunoResearch Laboratories) then streptavidin HRP. Slides were then stained with Opal™620 [9], counterstained with DAPI and mounted in Citifluor.

Samples were imaged on a Nikon A1R laser scanning confocal microscope. Image processing and γH2AX foci within $DAPI^+$ nuclei were enumerated with FIJI software. Pooled results from two independent experiments were reported.

#### Human samples

BM samples were obtained from MDS and normal patients in accordance with protocols approved by the Institutional ethics review boards as described [10,11]. $CD34^+$ cells were enriched from human BM samples and cDNA generated as previously described [10-12]. Somatic mutations were identified as previously described [11,12]. Patient sample details are listed in Table S5 and in [12].

#### HOXA1/PBX1/DNA model

The HOXA1-FL/PBX1/DNA model was created by threading the HOXA1-FL sequence (amino acids 202-290) onto the structure of HOXB1 in complex with DNA and PBX1 (PDB code: 1b72). The remaining 201 amino acids of the N-terminus are predicted to be unstructured. A model was also constructed using AlphaFold [13] as an independent confirmation of the overall structure.

A molecular dynamics simulation was carried out in GROMACS [14] (2019.6 release), using the AMBER99SB-ILDN forcefield [15] and the TIP3P water model [16]. The complex was positioned in a dodecahedral water box with 10 Å padding on all sides. Sodium and chloride ions were added to neutralize the charge of the system, and the system was energy minimized using the steepest descent algorithm until the maximum force in the system was less than 500 kJ/mol/nm. A subsequent NVT equilibration was performed for 100 ps at 310 K before a 10 ns production run conducted at 310 K.

**Immunoblot**

Western blot was performed on GP+E-86 ecotropic packaging cell lines overexpressing *Hoxa1* constructs. Nuclear and cytoplasm protein were isolated using the NE-PER® Nuclear and Cytoplasmic Extraction Reagents (Thermo Fisher) as per the manufacturer's instructions. A rabbit polyclonal anti-HOXA-1 antibody (Abcam; Cat No: ab64941) was used to detect HOXA1-FL. Antibodies detecting 53BP1 (Novus Biologicals, NB100-304) and pan actin (Thermo Scientific, MS-1295-P0) were used as loading controls.

**Whole Exome Sequencing**

Genomic DNA was isolated from whole bone marrow samples using the Puregene Cell Kit (QIAGEN), according to manufacturer's protocol. Samples from a minimum of 6 animals for each group were used for analyses. Libraries were prepared by GENEWIZ using the Agilent SureSelect$^{XT}$ Mouse All Exon system with sequencing performed on the Illumina Novaseq platform using 150 bp PE reads.

**Bioinformatic analysis**

Variant calling was done by GENEWIZ (GENEWIZ Ltd, Hong Kong). Cutadapt was applied to remove primers, adapters and low quality reads (Quality value < 20, over 10% N base reads, and reads shorter than 75 bp after trimming). For these analyses, the reference genome used was MGSCv37 (mm9), which was based on the C57BL/6J mouse strain.

Mutational landscapes were visualized using heatmaps generated in R (version ≥4.2) using the ComplexHeatmap package.

**Morphology**

PB smears (1-2 slides/mouse) were stained using May Grünwald/Giemsa (Amber Scientific). 5 μm sections of paraffin-embedded organs and decalcified bones were cut and stained with H & E. Morphological analyses were performed by two assessors independently (S.T. and M.W.). Representative micrographs were taken using the Microsystems DM 2000 microscope (Leica Microsystems) with a live camera interfaced to CellSens software (Olympus).

**Supplemental tables and figure legends**

Table S1. Peripheral blood parameters of the primary recipient mice at 16 weeks post-BMT.

| PB Parameters | MXIE | *WT-Hoxa1* | *MUT-Hoxa1* | *Hoxa1-T* |
|---|---|---|---|---|
| Leukocyte count ($x10^6$/ml) | 12 ± 5 | 9.75 ± 3.26 | 8.50 ± 2.88 * | 8.25 ± 5.88 * |
| Erythrocyte count ($x10^9$/ml) | 10.20 ± 0.70 | 10.15 ± 0.45 | 10.0 ± 1.07 | 9.65 ± 0.89 |
| Platelet count ($x10^6$/ml) | 674 ± 37 | 529 ± 40 | 447 ± 40 *** | 658 ± 57 |
| Hemoglobin (g/L) | 148 ± 9 | 145 ± 9 | 145 ± 14 | 143 ± 14 |
| MCV (fL) | 47.9 ± 1.70 | 47.6 ± 1.6 | 47.35 ± 2.25 | 48.25 ± 3.10 |

Shown are PB leukocyte counts, erythrocyte counts, platelet counts, hemoglobin and mean corpuscular volume (MCV) in *WT-Hoxa1, MUT-Hoxa1, Hoxa1-T* and MXIE overexpressing primary recipient mice at 16 weeks post transplantation. Leukocyte, Erythrocyte, Hemoglobin and MCV data are shown as median ± IQR, Platelet data are shown as mean ± SEM, n = 12-20 recipients. $^*p<0.05$, $^{***}p<0.001$ vs. MXIE (one-way ANOVA with multiple comparisons).

Table S2. Genes adjacent to retroviral integration sites of sAML mice.

| *WT-Hoxa1* | *MUT-Hoxa1* |
|---|---|
| *Meis1* | *Meis1* |
| *Meis1* | *Vpp1* |
| | *Sipa1l3* |
| | *Wdr87l* |
| | *Catsperg2* |
| | *MDS1 and EVI1 complex locus* |
| | *Fam187b* |
| | *Fxyd5* |
| | *Tha1* |
| | *Socs3* |

List of retroviral integration sites identified in sAML mice.

Table S3. Log-2 fold-change values of genes commonly altered in between *MUT-Hoxa1* MEPs and *WT-Hoxa1* MEPs compared to MXIE control MEPs.

| Gene | LogFC (MXIE vs *WT-Hoxa1*) | LogFC (MXIE vs *MUT-Hoxa1*) |
|---|---|---|
| *Tspan8* | -1.56 | -2.96 |
| *Ercc6l* | -1.22 | -1.71 |
| *Aldh1a1* | -1.20 | -1.69 |
| *Mt2* | -1.18 | -2.53 |
| *Rrm2* | -1.07 | -1.78 |
| *D930014E17Rik* | -1.05 | -1.14 |
| *Ass1* | -1.01 | -1.25 |
| *Cd2ap* | -1.00 | -1.31 |
| *Tra2a* | 1.09 | 1.45 |
| *Jun* | 1.15 | 1.57 |
| *Snora31* | 1.16 | 1.37 |
| *Lcp2* | 1.16 | 1.45 |
| *Zfp36* | 1.17 | 1.86 |
| *Gm614* | 1.18 | 1.94 |
| *ENSMUST00000129924* | 1.21 | 2.16 |
| *Vim* | 1.25 | 1.82 |
| *Tnfaip3* | 1.25 | 1.92 |
| *Slc18a2* | 1.27 | 1.48 |
| *Eif4e3* | 1.29 | 1.75 |
| *Eif4a2* | 1.30 | 1.24 |
| *Fosb* | 1.31 | 1.87 |
| *Gnb4* | 1.35 | 1.37 |
| *Fnbp1l* | 1.36 | 1.65 |
| *Adrbk2* | 1.36 | 1.57 |
| *Egr1* | 1.37 | 2.01 |
| *Ler2* | 1.38 | 2.11 |
| *Snora70* | 1.39 | 1.98 |
| *Srgap3* | 1.41 | 1.76 |
| *Ddx26b* | 1.42 | 2.41 |
| *Mir142* | 1.44 | 1.69 |
| *Phlda1* | 1.45 | 2.35 |
| *Gm10835* | 1.49 | 1.85 |
| *Klhl5* | 1.50 | 2.16 |
| *Rab44* | 1.64 | 2.06 |
| *10497248* | 1.68 | 1.61 |
| *Btg2* | 1.71 | 2.04 |
| *Snord14c* | 1.72 | 1.21 |
| *Lat* | 1.75 | 2.25 |
| *St8sia6* | 1.85 | 2.41 |
| *I730030J21Rik* | 2.05 | 2.23 |
| *Egr2* | 2.10 | 3.26 |
| *Cst7* | 2.14 | 2.63 |
| *Olfr248* | 2.31 | 2.82 |

| | | |
|---|---|---|
| *Scin* | 3.15 | 3.37 |
| *Tmem156* | 4.21 | 4.89 |
| *BC051162* | 4.30 | 4.91 |
| *Gpr65* | 4.55 | 4.92 |
| *Cpa3* | 4.60 | 4.64 |
| *Hoxa1* | 4.69 | 4.96 |
| *BC005685* | 4.73 | 5.46 |

List of genes and their Log-2 FC values commonly altered between *MUT-Hoxa1* MEPs and *WT-Hoxa1* MEPs compared to MXIE control MEPs.

Table S4. Expression levels of *HOXA* cluster transcripts in MDS patient BM $CD34^+$ cells.

| Gene | Fold increase | p value |
|---|---|---|
| *HOXA1* | 1.86 | 0.005 |
| *HOXA2* | 1.04 | 0.87 |
| *HOXA3* | 1.56 | 0.05 |
| *HOXA4* | 1.12 | 0.04 |
| *HOXA5* | 1.49 | 0.08 |
| *HOXA6* | 1.19 | 0.05 |
| *HOXA7* | 1.24 | 0.03 |
| *HOXA9* | 1.18 | 0.21 |
| *HOXA10* | 1.06 | 0.65 |
| *HOXA11* | 1 | 0.93 |
| *HOXA13* | 1 | 0.79 |

Fold-increase of *HOXA* cluster gene expression levels in BM $CD34^+$ from healthy and MDS patients obtained from a published gene expression profiling study [10] (183 MDS patients, 17 healthy controls).

Table S5. Summary of MDS patients analyzed for *HOXA1* isoform mRNA expression.

| Patient | Age | Disease type | Blast (%) | IPSS | IPSS-R |
|---|---|---|---|---|---|
| BM CD34$^+$ samples | | | | | |
| MDS1 | 35 | RAEB-I | 8 | 0.5 | 3 (low) |
| MDS2 | 73 | RAEB-I | 8 | 0.5 | 4 (intermediate) |
| MDS3 | 65 | RAEB-I | 9 | 1 | 6 (high) |
| MDS4 | 69 | RAEB-I | 6 | 1 | 4 (intermediate) |
| MDS5 | 41 | RAEB-I | 6 | 2 | 6.5 (very high) |
| MDS6 | 79 | RAEB-I | 9 | 1 | 4 (intermediate) |
| MDS7 | 66.7 | RAEB-I | 7 | 1 | 4 (intermediate) |
| MDS8 | 80.4 | RAEB-I | 5 | 2 | 7 (very high) |
| MDS9 | 59.8 | RAEB-I | 8 | 0.5 | 3 (low) |
| MDS10 | 68.1 | RAEB-I | 6 | 1.5 | 6.5 (very high) |
| MDS11 | 85 | RAEB-II | 15 | 1.5 | 5 (high) |
| MDS12 | 72 | RAEB-II | 14 | 1.5 | 5 (high) |
| MDS13 | 39 | RAEB-II | 10 | 1 | 5 (high) |
| MDS14 | 77 | RAEB-II | 17 | 1.5 | 5 (high) |
| MDS15 | 70 | RAEB-II | 13 | 3 | 8 (very high) |
| MDS16 | 69 | RAEB-II | 19 | 2.5 | 7.5 (very high) |
| MDS17 | 72 | RAEB-II | 15 | 2 | 5 (high) |
| MDS18 | 77 | RAEB-II | 12 | 2 | 6 (high) |
| MDS19 | 80 | RAEB-II | 19 | 1.5 | 4 (intermediate) |
| MDS20 | 77 | RAEB-II | 15 | 2 | 5 (high) |
| MDS21 | 70 | RAEB-II | 5 | 0.5 | 4 (intermediate) |
| MDS22 | 66 | RAEB-II | 14 | 1.5 | 5 (high) |
| MDS23 | 79 | RAEB-II | 18 | 2.5 | 7.5 (very high) |
| MDS24 | 67 | RAEB-II | 10 | 1 | 4 (intermediate) |
| MDS25 | 79 | RAEB-II | 19 | 2 | 6.5 (very high) |
| MDS26 | 78 | RAEB-II | 15 | 1.5 | 4.5 (intermediate) |
| MDS27 | 65 | RAEB-II | 10 | 0.5 | 3 (low) |
| MDS28 | 75 | RAEB-II | 12 | 1.5 | 3.5 (intermediate) |
| MDS29 | 70 | RAEB-II | 11 | 1.5 | 4.5 (intermediate) |
| MDS30 | 69.5 | RAEB-II | 11 | 1.5 | 4 (intermediate) |
| MDS31 | 72.1 | RAEB-II | 12 | 2 | 5.5 (high) |
| MDS32 | 68.7 | RAEB-II | 15 | 3 | 9 (very high) |
| MDS33 | 56.7 | RAEB-II | 15 | 3 | 8.5 (very high) |
| MDS34 | 71.4 | RAEB-II | 13 | 1.5 | 4.5 (intermediate) |
| MDS35 | 76.5 | RAEB-II | 13 | 2 | 7.5 (high) |

IPSS-R: Revised International Prognostic Scoring System

Patient information of MDS patients analyzed for *HOXA1* isoform expression in BM $CD34^+$ cells by qPCR.

Table S6. Lists of human and mouse QPCR primers and retroviral integration site analysis oligonucleotides.

Multiplexing QPCR primers

| Gene | Sequence |
|---|---|
| Murine *Hoxa1- FL* | 5' TTTACTCTGGAAACCTCTC 3' Forward Primer |
| | 5' CTGCTCTTGTCCATATGA 3' Reverse Primer |
| | 5' CATCACCACCACCACCAGG 3' Probe |
| Murine *Hoxa1- T* | 5' TATGGCCCCTATGGATTA 3' Forward Primer |
| | 5' AGGGTTTCTTTTAACTTTCAT 3' Reverse Primer |
| | 5' AGGAGGCAGACCCACCAAGAA 3' Probe |
| Murine *β2m* | 5' CAATAGTTGATCATATGCCAA 3' Forward Primer |
| | 5' TTACCAAAAGGAAAGTATGTC 3' Reverse Primer |
| | 5' CCTCTGTACTTCTCATTACTTGGATGC 3' Probe |
| Human *HOXA1-FL* | 5' TCGCCTCAATACATTCACCC 3' Forward Primer |
| | 5' TGGAGAGGGGACAAGGAGTT 3' Reverse Primer |
| Human *HOXA1-T* | 5' GAAGCAGACCCACCAAGAAG 3' Forward Primer |
| | 5' TTTGGGAGGGTTTCTTTTGA 3' Reverse Primer |
| Human *β2M* | 5' GTGGGATCGAGACATGTA 3' Forward Primer |
| | 5' GAGACAGCACTCAAAGTAGAA 3' Reverse Primer |

*β2m= β2-microglobulin*.

Retroviral integration site analysis oligonucleotides

| Oligos | Sequence |
|---|---|
| Splinkerette oligo # 1 | 5'CGAAGAGTAACCGTTGCTAGGAGAGACCGTGGCTGAATG AGACTGGTGTCGACACTAGTGG 3' |
| Splinkerette oligo # 2 | 5'GATCCCACTAGTGTCGACACCAGTCTCTAATTTTTTTTTTC AAAAAAA 3' |
| Long terminal repeat specific primer | 5' GCTAGCTTGCCAAACCTACAGGTGG 3' |
| Splinkerette specific primer | 5' CGAAGAGTAACCGTTGCTAGGAGAGACC 3' |
| Nested primer #1 | 5' GCCAAACCTACAGGTGGGGTCTTT 3' |
| Nested primer #2 | 5' GTGGCTGAATGAGACTGGTGTCGAC 3' |

Supplemental Dataset 1. Gene Expression Profiling data of MEPs and CMPs isolated from *MUT-Hoxa1*, *WT-Hoxa1* and MXIE primary recipients.

Supplemental Dataset 2. QuSAGE analyses of MEPs isolated from *MUT-Hoxa1*, *WT-Hoxa1* and MXIE primary recipients.

Supplemental Dataset 3. QuSAGE analyses of CMPs isolated from *MUT-Hoxa1*, *WT-Hoxa1* and MXIE primary recipients.

**Figure S1. Morphological evidence of leukemic infiltration in extramedullary tissues in sAML mice.**

Representative sections of mice shown in Figure 3: kidney, liver and spleen sections, respectively: (**A**, **D** and **G**) of a MXIE recipient control mouse, (**B**, **E** and **H**) of a *MUT-Hoxa1* recipient mouse that developed sAML (M1 subtype), and (**C**, **F** and **I**) of a *MUT-Hoxa1* recipient mouse that developed sAML (M6 subtype). Histological sections stained with hematoxylin and eosin showed tissue infiltration by blasts. Scale bar= 20µm.

**Figure S2. Microarray studies of MXIE, *WT-Hoxa1* and *MUT-Hoxa1* MEPs and CMPs.**

Shown are gene expression by MA plot with significant probesets [Abs(logFC) >1, Adj. P Value < 0.05] in red for MEPs: (**A)**, *MUT-Hoxa1* vs MXIE; (**B)**, *WT-Hoxa1* vs MXIE; for CMPs: (**C)**, *MUT-Hoxa1* vs MXIE; (**D)**, *WT-Hoxa1* vs MXIE.

**Figure S3. *HOXA1* isoform expression in human MDS**

Shown are (**A)**, Relative expression of *HOXA1-FL* and *HOXA1-T* in healthy controls and MDS patients classified according to IPSS. Lines connect the expression of *HOXA1-FL* and *HOXA1-T* in individual samples [23 healthy controls, 13 low-risk (Low), 13 intermediate-1 (Int-1), 9 Intermediate-2 (Int-2), 9 High-risk (High). *p<0.05; **p<0.01, ***p<0.001 vs. healthy controls, two-tailed paired Student's T-test. The somatic mutations with relative expression of *HOXA1-FL* and *HOXA1-T* for all 44 MDS patients are represented in (**B)**. The somatic mutations with *HOXA1-FL* and *HOXA1-T* expression (Q-

PCR) of the RAEB-1 and RAEB-2 patients shown in Figure 5D are represented in (**C**). IPSS and FAB classifications are shown for all samples.

**Figure S4. Hematopoietic parameters of *Hoxa1* knock-in BM.**

Shown are (**A**), schematic diagram of R26Cre$^{ERT2}$ *WT-Hoxa1*$^{ki/+}$, *MUT-Hoxa1*$^{ki/+}$ and Cre$^{+}$ control mice used to validate the expression of *Hoxa1-FL* and *Hoxa1-T* by Q-PCR in (**B**), cultured LKS+ and (**C**), cultured LKS- (n=4-6). (**D**) Heatmap showing gene-level mutations across individual donors stratified by genotype. The visualization highlights differences in pathway-specific mutation burden associated with genotype. Genes were grouped by functional category, displayed on the left of the heatmap, while sample-level annotations indicating mouse genotypes were displayed as top annotation bars. (**E**) Boxplot depicting the number of mutated genes per genotype. Data for B, C are shown as mean ± SEM, data for E are shown as median ± IQR. *p<0.05; *** p<0.001, ****p<0.0001 for comparisons between groups denoted by the bars. Data were analyzed using two-way ANOVA (B, C).

**Figure S5. Hematopoietic parameters of recipients of *Hoxa1* knock-in BM.**

Shown are analyses of transplant recipients of control, *WT- Hoxa1*$^{ki/+}$, *MUT- Hoxa1*$^{ki/+}$, *WT-Hoxa1*$^{ki/ki}$ and *MUT-Hoxa1*$^{ki/ki}$ BM at 15 months post-transplant. Shown are PB counts for: (**A**), leukocytes, (**B**), granulocytes, (**C**), B lymphocytes, (**D**), CD4+ T lymphocytes, (**E**) CD8$^{+}$ T lymphocytes. Numbers of cells in the BM: (**F**) Leukocytes, (**G**) immature granulocytes (Gr-1$^{low}$CD11b+), (**H**) mature granulocytes (Gr-1$^{br}$CD11b+), (**I**) CD4+ T lymphocytes, (**J**) CD8+ T lymphocytes. All data are shown as median ± IQR except (**D**), (**F**) and (**H**) which are mean ± SEM; *p<0.05; *** p<0.001 vs. Cre+ control (n=3-17) and analyzed by unpaired T-tests between mice with MDS and mice without MDS within a given genotype, Cre+ Control data are shown for visual comparisons. **(K)** Boxplot depicting the number of mutations comparing mice that developed MDS versus mice that did not develop MDS in each genotype. Data are shown as median ± IQR. **(L-M)** Heatmap showing gene-level mutation across individual recipients stratified by disease status and genotype. (**L**) Shows data for Cre + control and MDS recipients, (**M**) shows data for *Hoxa1* recipient mice that did not develop MDS. The visualization highlights differences in pathway-specific mutation burden associated with genotype and disease state. Genes were grouped by functional category, displayed on the left of the heatmap, while sample-level annotations indicating disease status and donor genotype were displayed as top annotation bars.

#### References


1. Purton LE, Bernstein ID, Collins SJ. All-trans retinoic acid delays the differentiation of primitive hematopoietic precursors (lin-c-kit+Sca-1(+)) while enhancing the terminal maturation of committed granulocyte/monocyte progenitors. *Blood*. 1999;94(2):483–495.
2. Purton LE, Dworkin S, Olsen GH, et al. RARgamma is critical for maintaining a balance between hematopoietic stem cell self-renewal and differentiation. *J Exp Med*. 2006;203(5):1283–1293.
3. Uren AG, Mikkers H, Kool J, et al. A high-throughput splinkerette-PCR method for the isolation and sequencing of retroviral insertion sites. *Nat Protoc*. 2009;4(5):789–798.
4. Irizarry RA, Bolstad BM, Collin F, Cope LM, Hobbs B, Speed TP. Summaries of Affymetrix GeneChip probe level data. *Nucleic Acids Res*. 2003;31(4):e15.
5. Carvalho BS, Irizarry RA. A framework for oligonucleotide microarray preprocessing. *Bioinformatics*. 2010;26(19):2363–2367.
6. Smyth GK. Linear models and empirical bayes methods for assessing differential expression in microarray experiments. *Stat Appl Genet Mol Biol*. 2004;3:Article3.
7. Yaari G, Bolen CR, Thakar J, Kleinstein SH. Quantitative set analysis for gene expression: a method to quantify gene set differential expression including gene-gene correlations. *Nucleic Acids Res*. 2013;41(18):e170.
8. Nakamura A, Sedelnikova OA, Redon C, et al. Techniques for gamma-H2AX detection. *Methods Enzymol*. 2006;409:236–250.
9. Green AC, Rudolph-Stringer V, Straszkowski L, et al. Retinoic Acid Receptor gamma Activity in Mesenchymal Stem Cells Regulates Endochondral Bone, Angiogenesis, and B Lymphopoiesis. *J Bone Miner Res*. 2018;33(12):2202–2213.

10. Pellagatti A, Cazzola M, Giagounidis A, et al. Deregulated gene expression pathways in myelodysplastic syndrome hematopoietic stem cells. *Leukemia*. 2010;24(4):756–764.
11. Unnikrishnan A, Papaemmanuil E, Beck D, et al. Integrative Genomics Identifies the Molecular Basis of Resistance to Azacitidine Therapy in Myelodysplastic Syndromes. *Cell Rep*. 2017;20(3):572–585.
12. Im H, Rao V, Sridhar K, et al. Distinct transcriptomic and exomic abnormalities within myelodysplastic syndrome marrow cells. *Leuk Lymphoma*. 2018;59(12):2952–2962.
13. Jumper J, Evans R, Pritzel A, et al. Highly accurate protein structure prediction with AlphaFold. *Nature*. 2021;596(7873):583–589.
14. Pronk S, Pall S, Schulz R, et al. GROMACS 4.5: a high-throughput and highly parallel open source molecular simulation toolkit. *Bioinformatics*. 2013;29(7):845–854.
15. Lindorff-Larsen K, Piana S, Palmo K, et al. Improved side-chain torsion potentials for the Amber ff99SB protein force field. *Proteins*. 2010;78(8):1950–1958.
16. Jorgensen WL, Chandrasekhar J, Madura JD, Impey RW, Klein ML. Comparison of Simple Potential Functions for Simulating Liquid Water. *Journal of Chemical Physics*. 1983;79:926–935.

Figure S1

#### Tissue infiltration by leukemic blasts

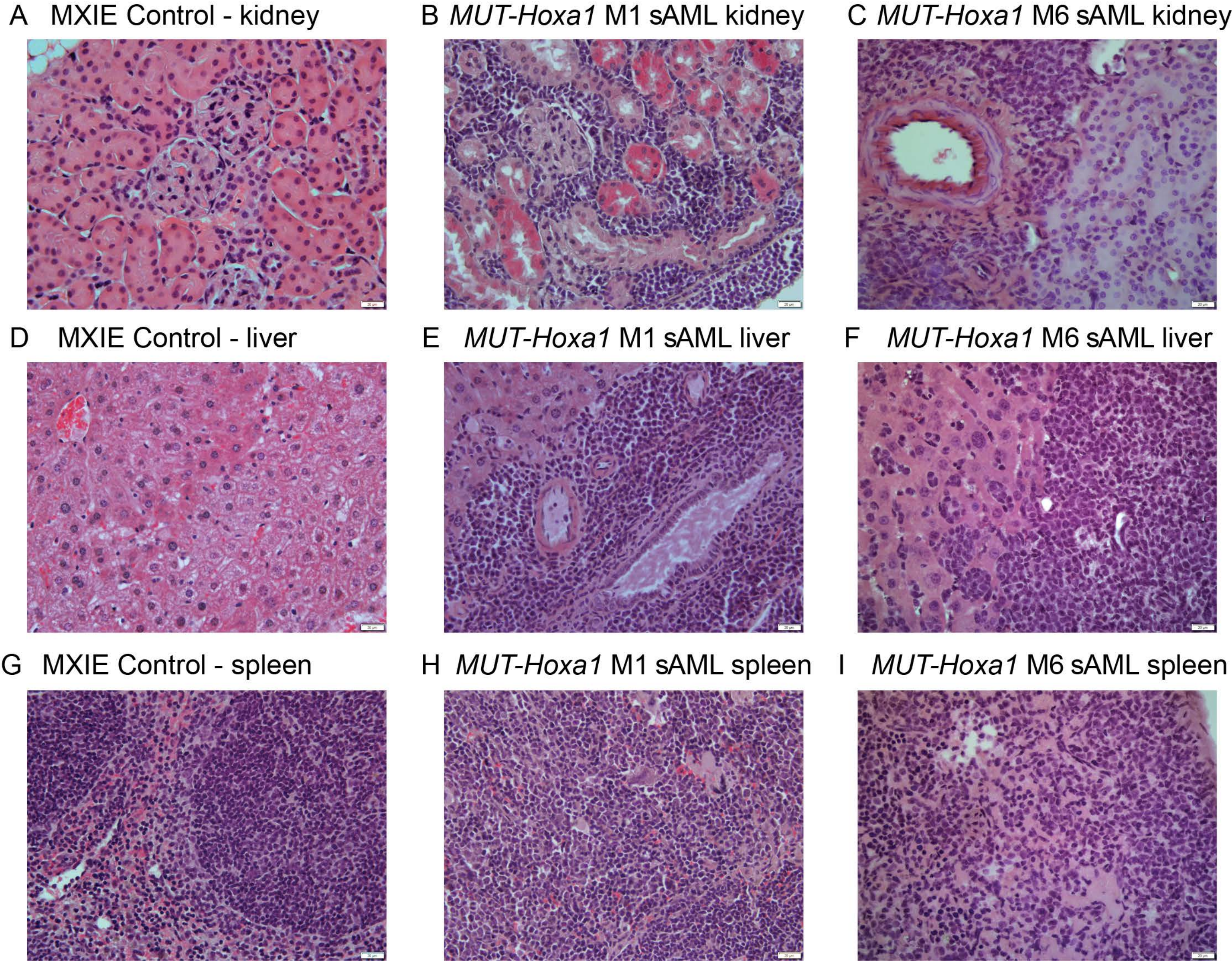

Figure S2

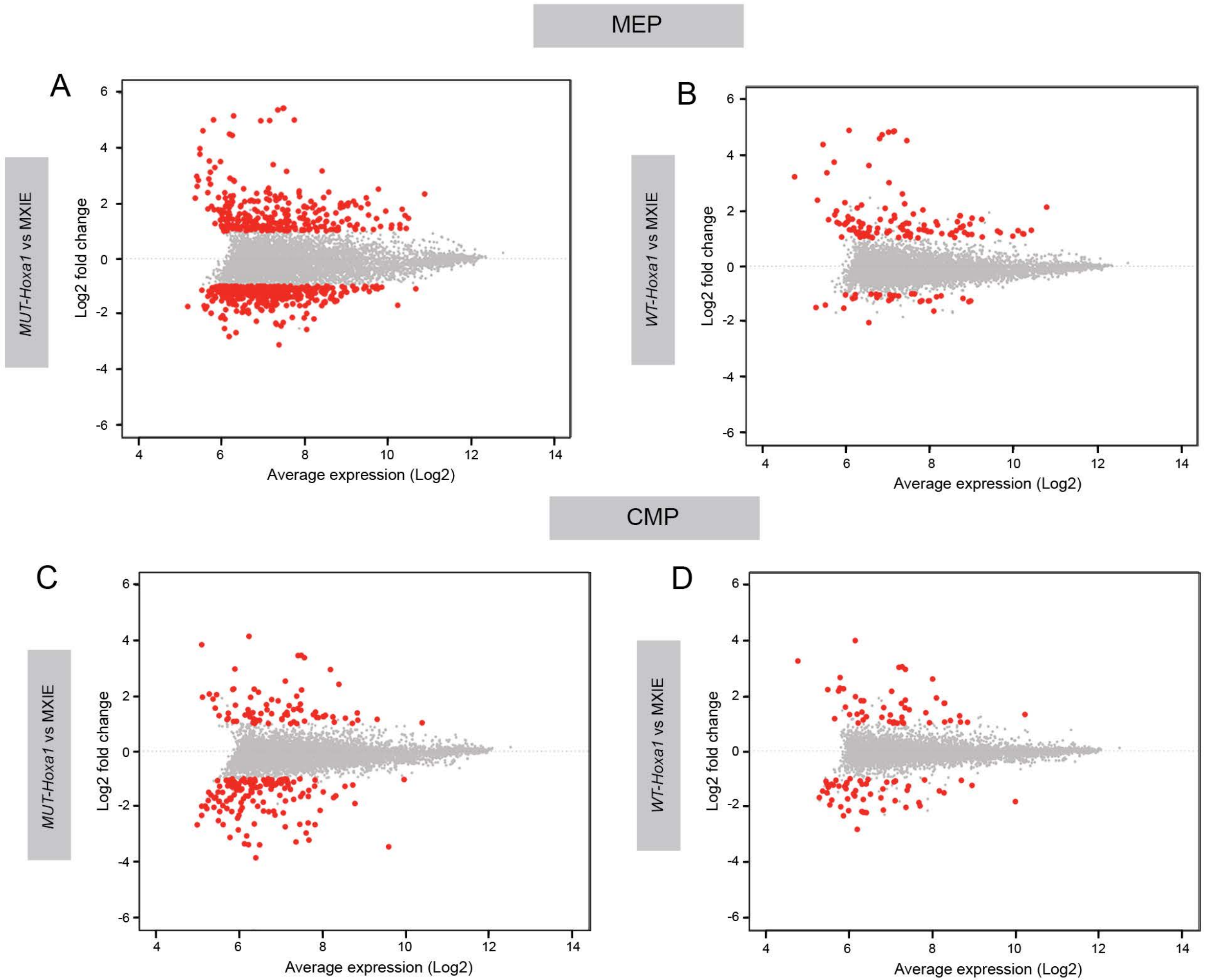
MEP
A
MUT-Hoxa1 vs MXIE
Log2 fold change
Average expression (Log2)
B
WT-Hoxa1 vs MXIE
Log2 fold change
Average expression (Log2)
CMP
C
MUT-Hoxa1 vs MXIE
Log2 fold change
Average expression (Log2)
D
WT-Hoxa1 vs MXIE
Log2 fold change
Average expression (Log2)

Figure S3

A

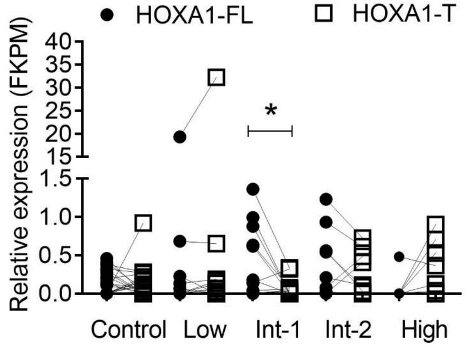

HOXA1-FL
HOXA1-T
Relative expression (FKPM)
40
35
30
25
20
15
1.5
1.0
0.5
0.0
*
Control
Low
Int-1
Int-2
High


B

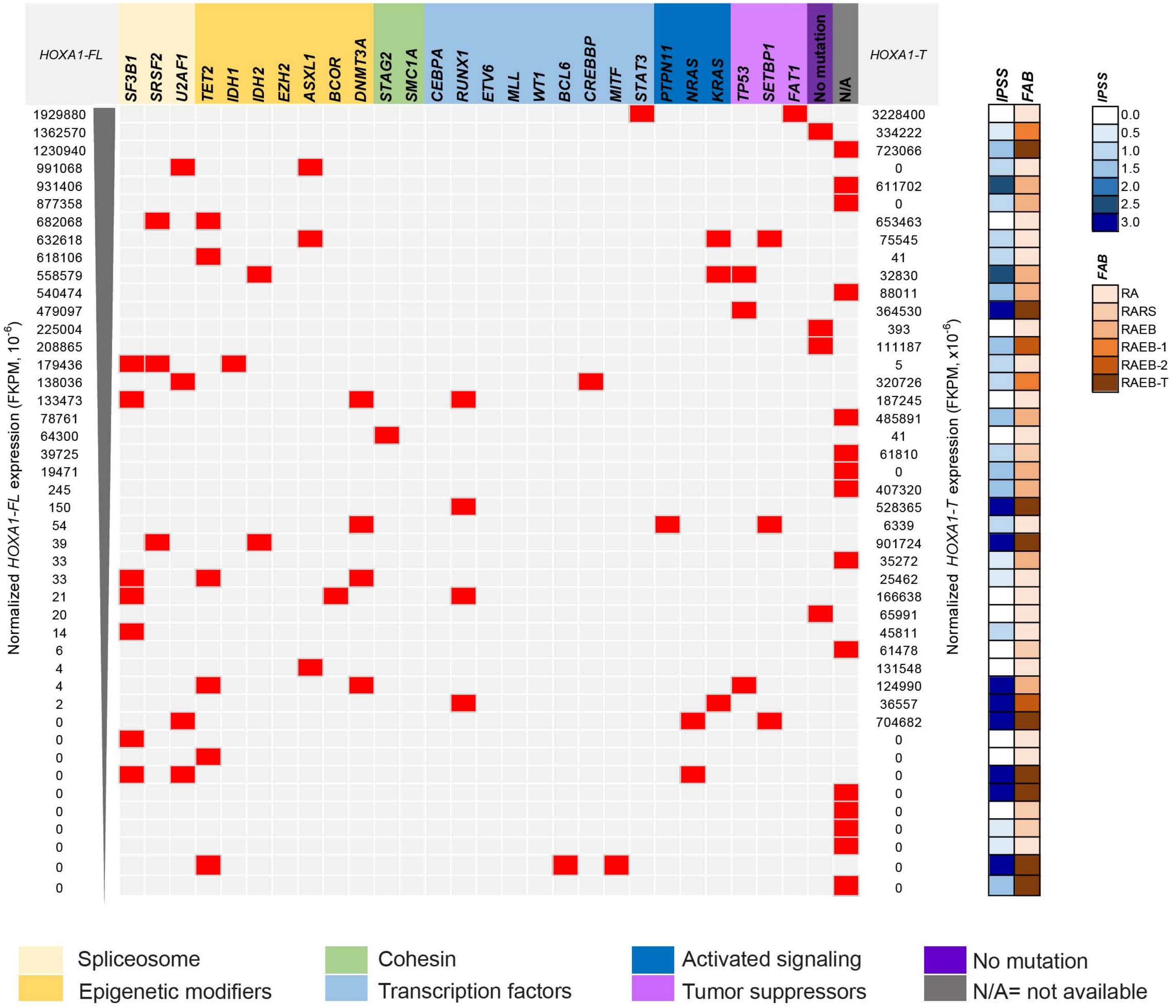

HOXA1-FL
SF3B1
SRSF2
U2AF1
TET2
IDH1
IDH2
EZH2
ASXL1
BCOR
DNMT3A
STAG2
SMC1A
CEBPA
RUNX1
ETV6
MLL
WT1
BCL6
CREBBP
MITF
STAT3
PTPN11
NRAS
KRAS
TP53
SETBP1
FAT1
No mutation
N/A
HOXA1-T
IPSS
FAB
Normalized HOXA1-FL expression (FKPM, $10^{-6}$)
1929880
1362570
1230940
991068
931406
877358
682068
632618
618106
558579
540474
479097
225004
208865
179436
138036
133473
78761
64300
39725
19471
245
150
54
39
33
33
21
20
14
6
4
4
2
0
0
0
0
0
0
0
0
0
0
Normalized HOXA1-T expression (FKPM, $x10^{-6}$)
3228400
334222
723066
0
611702
0
653463
75545
41
32830
88011
364530
393
111187
5
320726
187245
485891
41
61810
0
407320
528365
6339
901724
35272
25462
166638
65991
45811
61478
131548
124990
36557
704682
0
0
0
0
0
0
0
0
0
IPSS
0.0
0.5
1.0
1.5
2.0
2.5
3.0
FAB
RA
RARS
RAEB
RAEB-1
RAEB-2
RAEB-T
Spliceosome
Epigenetic modifiers
Cohesin
Transcription factors
Activated signaling
Tumor suppressors
No mutation
N/A= not available

Figure S3 continued

C

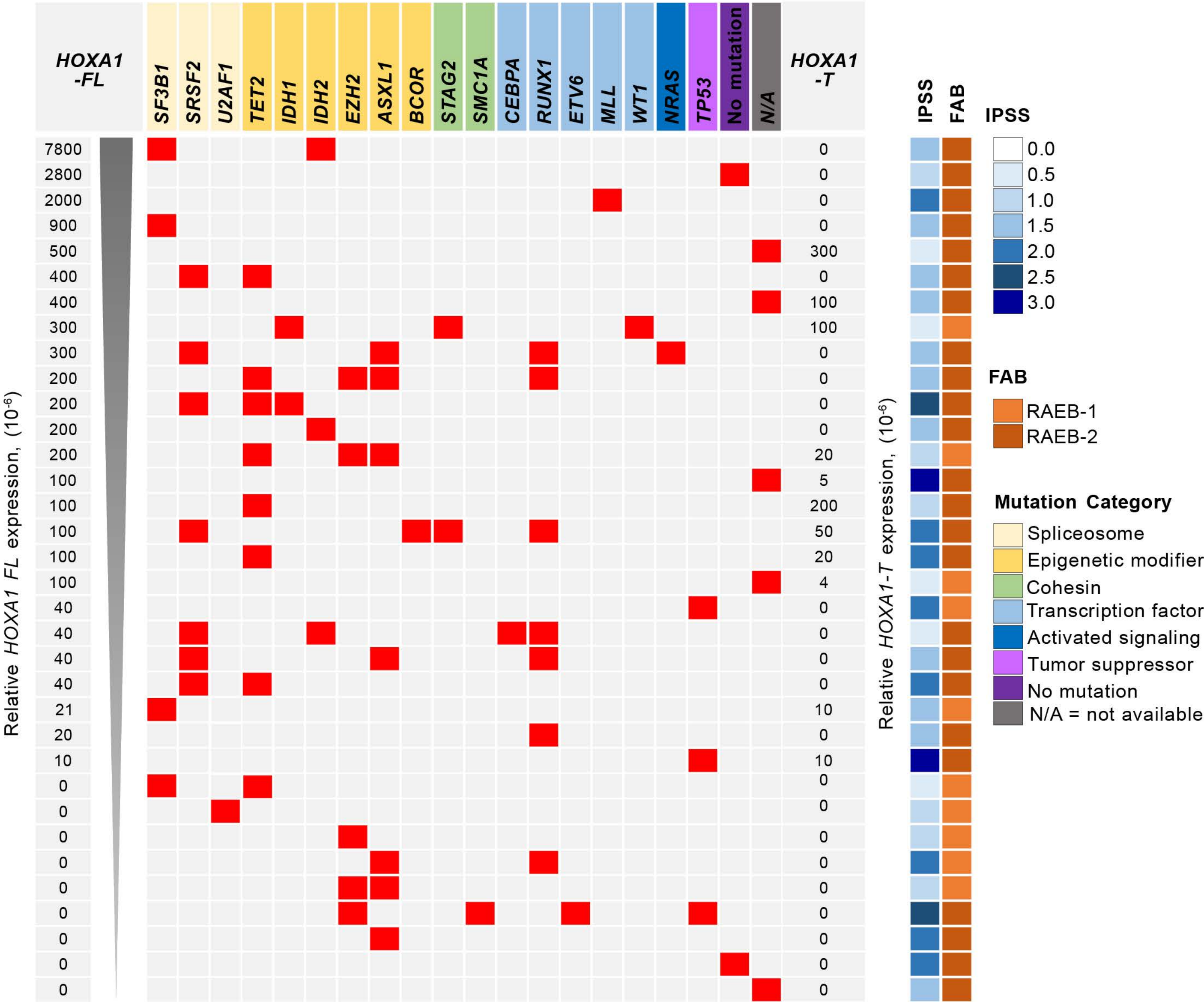

### Figure S4

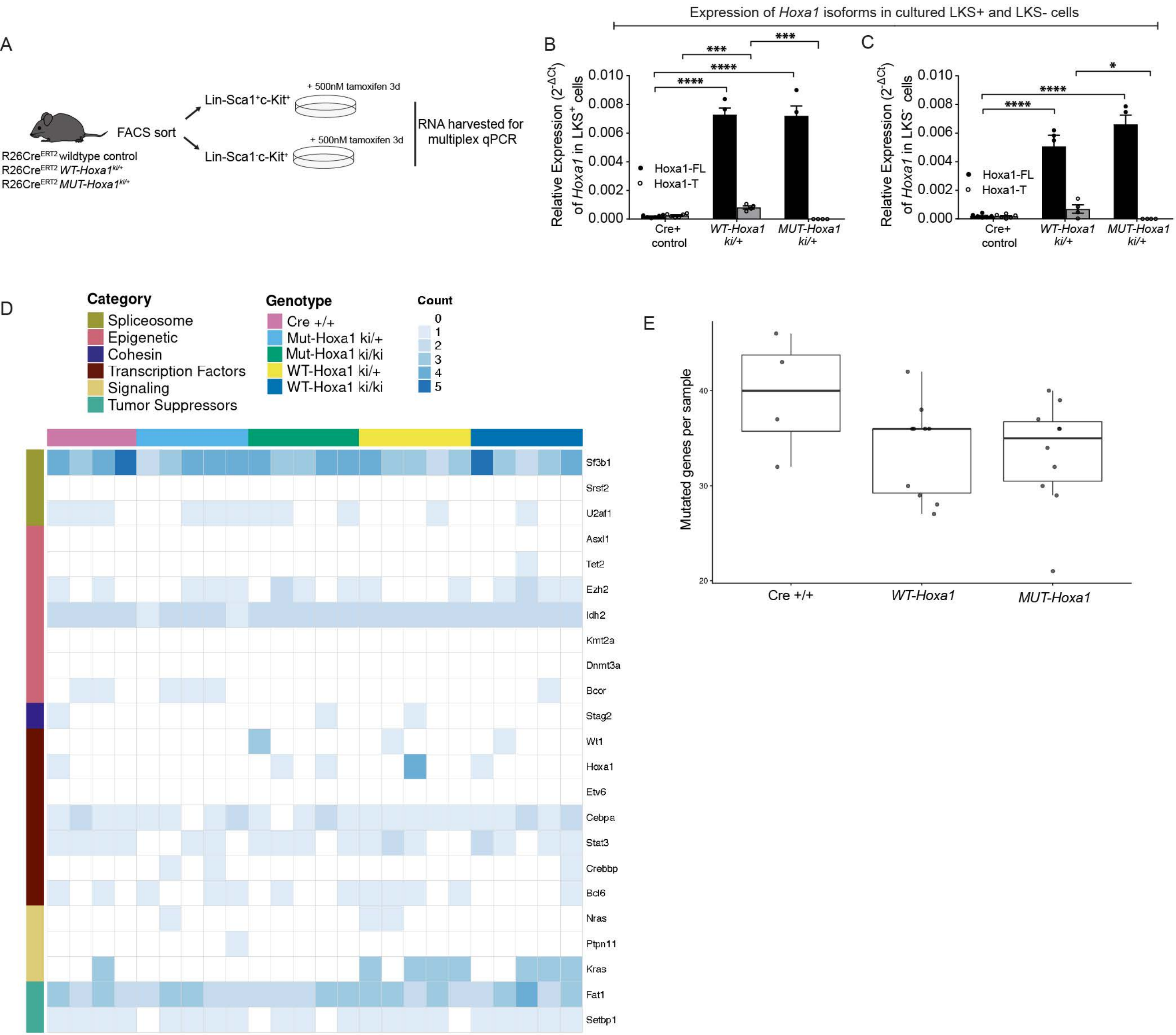
Expression of Hoxa1 isoforms in cultured LKS+ and LKS- cells
A
FACS sort
R26Cre^ERT2 wildtype control
R26Cre^ERT2 WT-Hoxa1 ki/+
R26Cre^ERT2 MUT-Hoxa1 ki/+
Lin-Sca1+c-Kit+
Lin-Sca1-c-Kit+
+ 500nM tamoxifen 3d
RNA harvested for multiplex qPCR
B
Relative Expression (2^-ΔCt) of Hoxa1 in LKS+ cells
Hoxa1-FL
Hoxa1-T
Cre+ control
WT-Hoxa1 ki/+
MUT-Hoxa1 ki/+
C
Relative Expression (2^-ΔCt) of Hoxa1 in LKS- cells
D
Category
Spliceosome
Epigenetic
Cohesin
Transcription Factors
Signaling
Tumor Suppressors
Genotype
Cre +/+
Mut-Hoxa1 ki/+
Mut-Hoxa1 ki/ki
WT-Hoxa1 ki/+
WT-Hoxa1 ki/ki
Count
Sf3b1
Srsf2
U2af1
Asxl1
Tet2
Ezh2
Idh2
Kmt2a
Dnmt3a
Bcor
Stag2
Wt1
Hoxa1
Etv6
Cebpa
Stat3
Crebbp
Bcl6
Nras
Ptpn11
Kras
Fat1
Setbp1
E
Mutated genes per sample
Cre +/+
WT-Hoxa1
MUT-Hoxa1

Figure S5

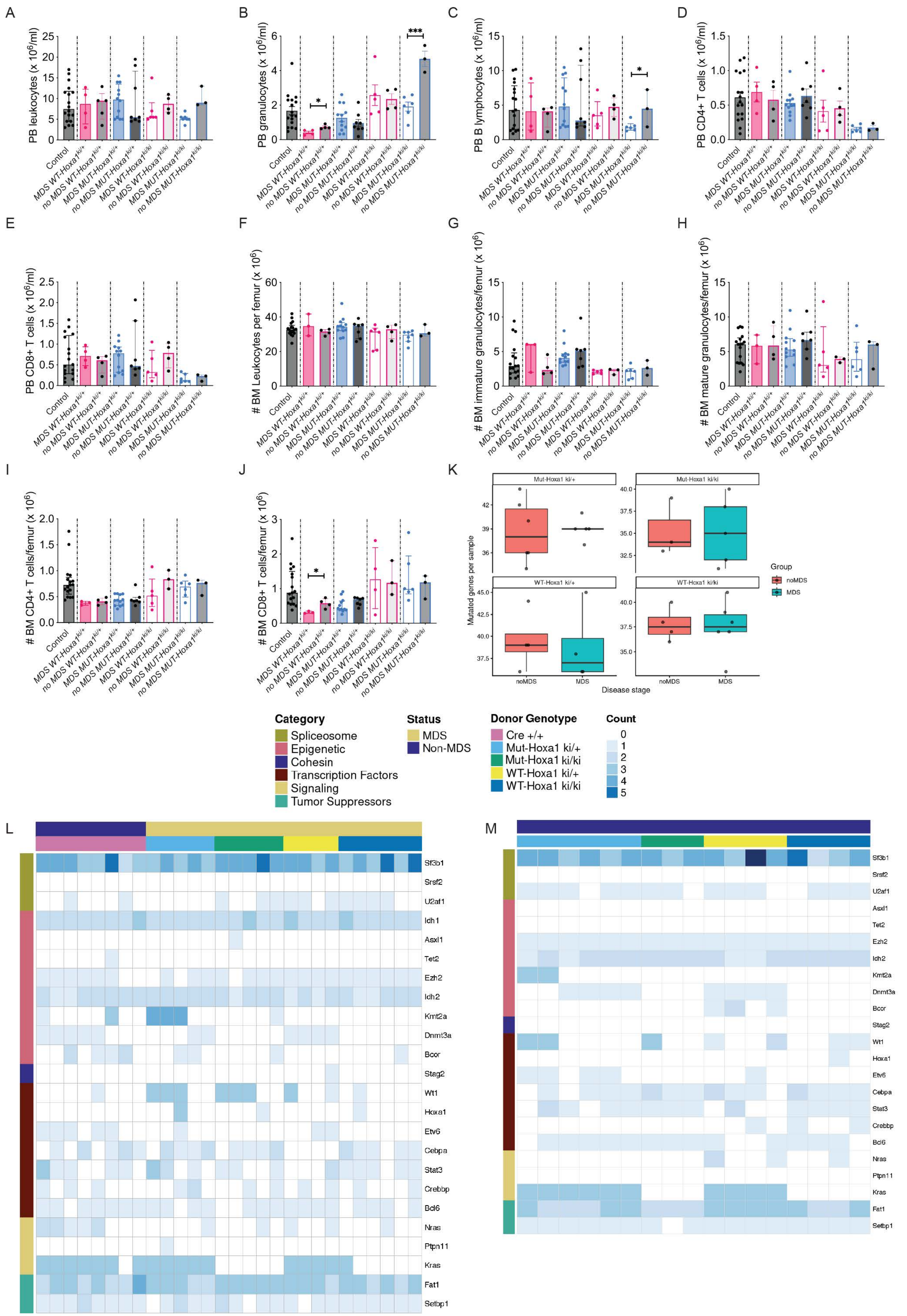

A
PB leukocytes (x 10^6/ml)
B
PB granulocytes (x 10^6/ml)
C
PB B lymphocytes (x 10^6/ml)
D
PB CD4+ T cells (x 10^6/ml)
E
PB CD8+ T cells (x 10^6/ml)
F
### BM Leukocytes per femur (x 10^6)
G
### BM immature granulocytes/femur (x 10^6)
H
### BM mature granulocytes/femur (x 10^6)
I
### BM CD4+ T cells/femur (x 10^6)
J
### BM CD8+ T cells/femur (x 10^6)
Control
MDS WT-Hoxa1ki/+
no MDS WT-Hoxa1ki/+
MDS MUT-Hoxa1ki/+
no MDS MUT-Hoxa1ki/+
MDS WT-Hoxa1ki/ki
no MDS WT-Hoxa1ki/ki
MDS MUT-Hoxa1ki/ki
no MDS MUT-Hoxa1ki/ki
*
***
K
Mut-Hoxa1 ki/+
Mut-Hoxa1 ki/ki
WT-Hoxa1 ki/+
WT-Hoxa1 ki/ki
Mutated genes per sample
Disease stage
noMDS
MDS
Group
noMDS
MDS
Category
Spliceosome
Epigenetic
Cohesin
Transcription Factors
Signaling
Tumor Suppressors
Status
MDS
Non-MDS
Donor Genotype
Cre +/+
Mut-Hoxa1 ki/+
Mut-Hoxa1 ki/ki
WT-Hoxa1 ki/+
WT-Hoxa1 ki/ki
Count
0
1
2
3
4
5
L
Sf3b1
Srsf2
U2af1
Idh1
Asxl1
Tet2
Ezh2
Idh2
Kmt2a
Dnmt3a
Bcor
Stag2
Wt1
Hoxa1
Etv6
Cebpa
Stat3
Crebbp
Bcl6
Nras
Ptpn11
Kras
Fat1
Setbp1
M
Sf3b1
Srsf2
U2af1
Asxl1
Tet2
Ezh2
Idh2
Kmt2a
Dnmt3a
Bcor
Stag2
Wt1
Hoxa1
Etv6
Cebpa
Stat3
Crebbp
Bcl6
Nras
Ptpn11
Kras
Fat1
Setbp1
